# The CTCF Paralog BORIS Contributes to the Ovarian Cancer Transcriptional Program by Relaxing CTCF-mediated 3D Genome Organization

**DOI:** 10.64898/2026.08.04.742596

**Authors:** Dharmendra Nath Bhatt, Arielle Scott, Ludmila Krymskaya, Tovah E. Markowitz, Emma Price, Yon Ji, Dmitri Loukinov, Maria Traver, Joseph Brzostowski, Victor V. Lobanenkov, Elena M. Pugacheva

## Abstract

Disruption of three-dimensional genome architecture is a major driver of cancer initiation and progression, frequently arising from genetic and epigenetic alterations at CTCF- and cohesin-bound chromatin anchors. These same chromatin loop anchors can also be occupied by the germ cell-specific CTCF paralog CTCFL (BORIS), which is aberrantly activated in multiple malignancies. Here, we show that in ovarian cancer cells, BORIS activation establishes a distinct transcriptional program that collapses following loss of BORIS chromatin binding and is accompanied by widespread changes in CTCF and cohesin occupancy, histone modifications, and chromatin accessibility. These BORIS-dependent transcriptional alterations occur in long-range genomic clusters, resulting in the coordinated activation or repression of neighboring genes, including hormonally regulated gene families. BORIS loss also increases topologically associating domain (TAD) insulation, strengthens A/B compartment segregation and chromatin loop interactions, and results in a more compact and constrained chromatin architecture. Together, our findings suggest that aberrant BORIS activation promotes transcriptional reprogramming by weakening CTCF-mediated chromatin insulation and relaxing three-dimensional genome organization in ovarian cancer.

**Significance:** High-grade serous ovarian carcinoma, the most common subtype of ovarian cancer, remains one of the deadliest gynecological malignancies because of its late diagnosis, extensive genomic instability, and frequent therapeutic resistance. The lack of reliable biomarkers for early detection underscores the need to identify new molecular drivers of disease initiation and progression. Here, we show that the germline-specific gene CTCFL (BORIS) is aberrantly activated in high-grade serous ovarian carcinoma, where it remodels transcriptional, epigenetic, and three-dimensional genome organization by occupying CTCF-bound chromatin loop anchors and weakening CTCF-mediated chromatin insulation. These findings identify BORIS as a key regulator of transcriptional reprogramming in ovarian cancer and support its further investigation as both a prognostic biomarker and a potential therapeutic target.

## Introduction

Malignant transformation of normal cells is driven by cancer-associated transcriptional reprogramming, which is often supported by aberrant activity of transcription factors and alterations in three-dimensional (3D) genome organization (1). Two key architectural proteins, CTCF and cohesin, are essential for maintaining 3D genome organization in vertebrate cells (2–4); yet they also exhibit cancer-specific binding alterations, gaining access to novel genomic sites or losing occupancy at loci normally bound in healthy tissues (5). These changes disrupt CTCF-mediated chromatin anchors and promote the formation of oncogenic chromatin loops and aberrant topologically associating domains (TADs) (6). The resulting reorganization of chromatin topology was shown to activate proto-oncogenes (7), silence tumor-suppressor genes (8), and dysregulate core transcriptional programs (9), thereby promoting cancer initiation and progression (1).

CTCF is a multifunctional architectural protein with an 11-zinc-finger (11ZF) DNA-binding domain (10). Through combinatorial use of its 11 ZFs, CTCF recognizes diverse DNA sequences (11) and halts cohesin-mediated loop extrusion via its N-terminal domain (12, 13), thereby establishing anchors for 3D genome organization. Aberrant CpG methylation (14), changes in histone modifications (15), altered nucleosome positioning (16), differential transcription factor occupancy (17), and mutations (18) or single-nucleotide polymorphisms (SNPs) within CTCF consensus motifs (19) can lead to the loss or gain of CTCF binding sites. These alterations disrupt local and global chromatin architecture, contributing to the 3D genome reorganization observed in many disease states, including cancer (20).

Another factor that may drive the reorganization of CTCF-mediated chromatin loops in cancer is CTCFL (also known as BORIS, Brother of the Regulator of Imprinted Sites), a paralog of CTCF that is normally restricted to gametogenesis but aberrantly reactivated in multiple malignancies (21). Similar to CTCF, BORIS contains an almost identical 11-zinc finger (11ZF) DNA-binding domain, allowing it to recognize the same DNA motifs (22). However, the N- and C-terminal domains of BORIS differ substantially from CTCF’s termini (22). Unlike CTCF, the BORIS N-terminus does not interact with the cohesin complex in somatic cells and therefore cannot halt cohesin-mediated loop extrusion (12, 13). Moreover, BORIS lacks intrinsic pioneer activity and depends on other chromatin factors for DNA accessibility, whereas CTCF can independently open chromatin to establish new binding sites (15, 23). Experimental evidence from endogenous and ectopic expression models indicates that BORIS prefers to bind with CTCF at clustered CTCF-binding regions (2×CTCF sites, defined by two or more adjacent CTCF motifs) (24–26). Such interactions may alter CTCF-binding dynamics and reshape local chromatin topology, leading to transcriptional reprogramming in cancer cells. Consistent with this notion, BORIS has been implicated in the reorganization of chromatin loops that drive resistance to anaplastic lymphoma kinase (ALK) inhibitors in MYCN-amplified neuroblastoma (27).

During spermatogenesis, CTCF and BORIS co-occupy clustered CTCF-binding sites to regulate testis-specific gene expression programs (25, 26). When aberrantly reactivated in cancer, BORIS can similarly interact with CTCF, partially recapitulating their cooperative functions observed in germ cells (24, 25, 28, 29). In somatic contexts, BORIS co-binding with CTCF has been shown to remodel CTCF-binding sites into active promoters (24), thereby driving the ectopic activation of testis-specific, or so-called cancer-testis genes. This interaction may also rewire CTCF-mediated chromatin loops and alter higher-order genome topology; however, this possibility has not yet been systematically investigated.

Recent studies have proposed testis-specific BORIS as a prognostic and therapeutic marker for ovarian cancers (30–32). Despite the high frequency of aberrant BORIS activation in ovarian tumors, its precise functional role remains elusive. We previously demonstrated that ectopic BORIS expression in BORIS-negative fallopian tube cells increases chromatin accessibility, stabilizes CTCF occupancy, and activates genes involved in cell motility and invasion (30). In the present study, we generated a knockout of BORIS in BORIS-positive ovarian cancer cells. We demonstrate that BORIS loss causes widespread transcriptional deregulation, accompanied by both the loss and gain of CTCF binding sites, epigenetic alterations, and global rewiring of CTCF-mediated chromatin loops, leading to profound changes in gene expression. Taken together, our findings indicate that aberrant BORIS activation may profoundly reshape 3D genome architecture in cancer cells through modulation of CTCF binding and chromatin organization.

## Results

### BORIS/CTCFL is a germline-specific factor aberrantly activated in Müllerian-derived cancers

As a definitive Cancer-Testis Antigen (CTA), BORIS aberrant activation and expression was described in many primary cancers and cell lines (21, 22). To systematically evaluate BORIS expression across human cancers, we analyzed multiple publicly available RNA-seq datasets comprising primary tumor samples from diverse cancer types. Analysis of GEPIA (Gene Expression Profiling Interactive Analysis) data (33) revealed that BORIS expression was elevated in tumors compared with normal tissues across many types of cancer, although the extent of upregulation varied considerably. Among the cancers analyzed, CESC (cervical squamous cell carcinoma and endocervical adenocarcinoma), OV (ovarian serous cystadenocarcinoma), UCEC (uterine corpus endometrial carcinoma), and UCS (uterine carcinosarcoma) exhibited particularly high and significant BORIS expression (Fig. 1A). BORIS activation was also frequently associated with amplification of the BORIS locus and heterozygous deletion of its paralog, CTCF (SI Appendix, Fig. S1A).

**Figure 1.**
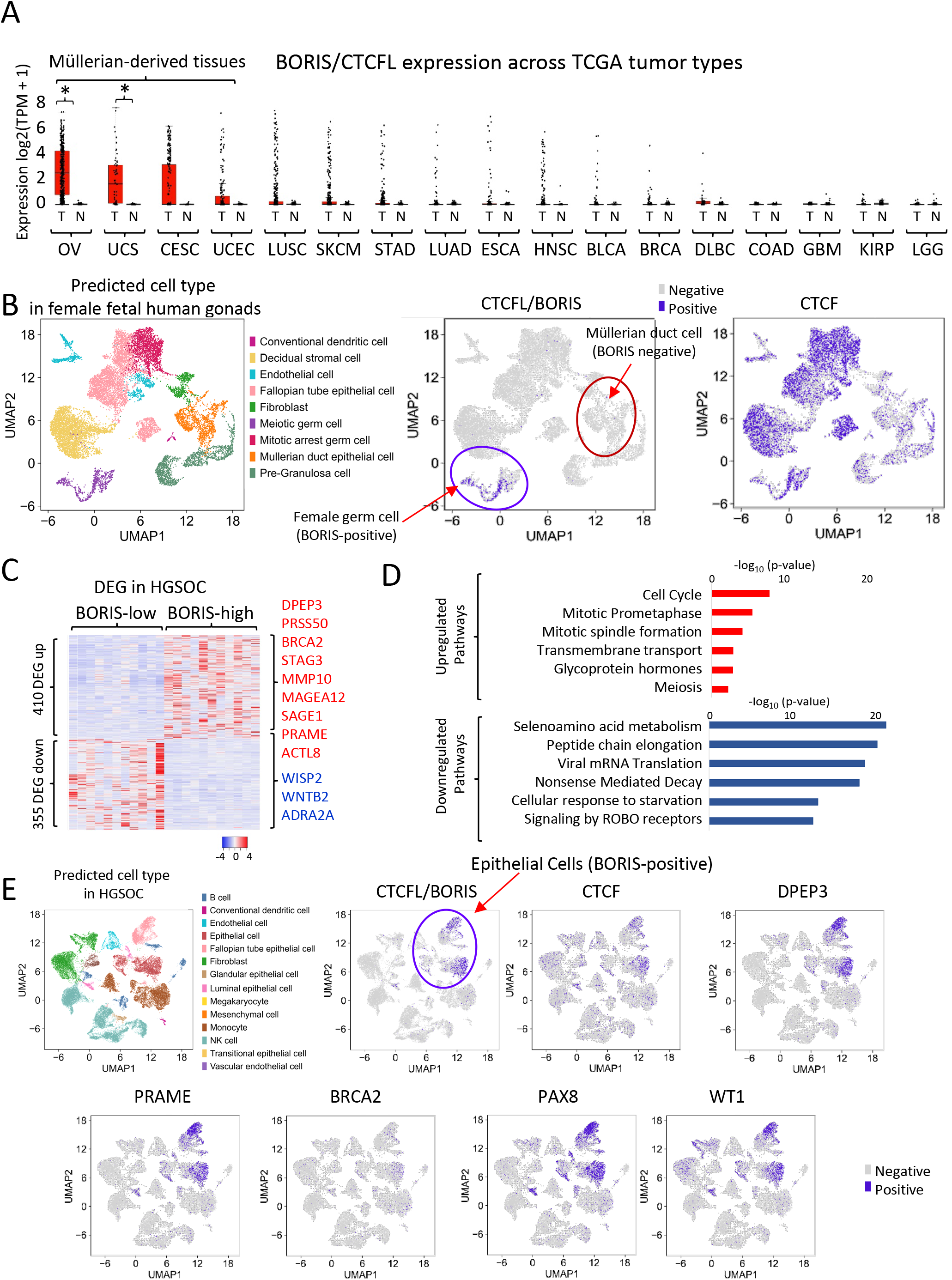
BORIS is aberrantly expressed in ovarian cancer and is associated with a germ cell-like transcriptional program. **(A)** Expression of CTCFL/BORIS in different types of cancer: comparisons of tumors (T) with the normal counterparts (N). OV (Ovarian Cancer: number of tumor samples=426, number of normal samples=88), UCS (Uterine carcinosarcoma: T=57, N=78), CESC (Cervical Squamous Cell Carcinoma and Endocervical Adenocarcinoma: T=306, N=13), UCEC (Uterine Corpus Endometrial Carcinoma: T=174, N=91), LUSC (Lung Squamous Cell Carcinoma: T=486, N=338), SKCM (Skin Cutaneous Melanoma: T=461, N=558), STAD (Stomach adenocarcinoma: T=408, N=211), LUAD (Lung Adenocarcinoma: T=483, N=347), ESCA (Esophageal carcinoma: T=182, N=286), HNSC (Head and Neck Squamous Cell Carcinoma: T=519, N=44), BLCA (Bladder Cancer: T=404, N=28), BRCA (Breast cancer: T=1085, N=291), DLBC (Diffuse Large B-cell Lymphoma: T=47, N=337), COAD (Colon Adenocarcinoma: T=275, N=349), GBM (Glioblastoma: T=163, N=207), KIRP (Kidney Renal Papillary Cell Carcinoma: T=286, N=60), LGG (Low-Grade Glioma: T=518, N=207). *p < 0.01. **(B)** Single-cell RNA-seq analysis of the developing female reproductive tract. UMAP projections show annotated cell populations (left) and expression of *BORIS/CTCFL* (middle) and *CTCF* (right). *BORIS* expression is restricted to female germ cells, whereas *CTCF* is expressed across all cell types. **(C)** Heatmap of the 765 genes differentially expressed between BORIS-low and BORIS-high high-grade serous ovarian carcinoma (HGSOC) tumors. Selected BORIS-associated genes are indicated. **(D)** Gene ontology/pathway enrichment analysis of genes upregulated (red) or downregulated (blue) in BORIS-high HGSOC tumors. **(E)** Single-cell RNA-seq analysis of HGSOC tumors. UMAP projections show annotated cell populations (left) and expression of BORIS/CTCFL, CTCF, WT1, DPEP3, PRAME, and BRCA2, PAX8, WT1. Expression of BORIS overlaps with epithelial tumor cells expressing ovarian cancer markers (red arrow).

Interestingly, the cancers with the highest BORIS expression are all gynecologic malignancies arising from the female reproductive tract. These cancers share several notable features. First, they originate from Müllerian-derived tissues, which develop from the embryonic Müllerian ducts and give rise to the fallopian tubes, uterus, cervix, and upper vagina (34). Second, they are generally hormone-responsive tumors (35). Third, they frequently harbor alterations in the PI3K/AKT/mTOR signaling pathway (36). The enrichment of BORIS expression in these related malignancies suggests that common developmental and regulatory programs may contribute to its activation.

The expression of BORIS in Müllerian-derived cancers is unexpected, as BORIS has been considered a testis-specific factor expressed predominantly in male germ cells, with highest levels in spermatogonia (26). To determine whether BORIS is also expressed in female cells under physiological conditions, we analyzed published single-cell RNA-sequencing (scRNA-seq) datasets from human female embryonic tissues, including germ cells undergoing meiotic entry, as well as developing Müllerian duct cells (37). We detected no BORIS expression in embryonic Müllerian-derived cells (Fig. 1B). In contrast, BORIS expression was readily detected in primordial germ cells and oogonia during the first and second trimesters of female fetal development (Fig. 1B). These findings demonstrate that BORIS is expressed in both male and female germ cells, establishing BORIS as a germline-specific rather than a male germ cell– specific protein. Moreover, the absence of BORIS expression in normal Müllerian-derived cells indicates that its activation in ovarian cancer does not reflect persistence of a Müllerian developmental program but rather ectopic reactivation of a germline-specific transcriptional regulator.

Analysis of RNA-seq data from 300 primary high-grade serous ovarian carcinomas (HGSOCs) in The Cancer Genome Atlas (TCGA) (38) revealed detectable BORIS expression in 86% of tumors (SI Appendix, Fig. S1B,C). Despite its widespread expression, BORIS levels varied markedly among patients, highlighting substantial heterogeneity in BORIS activation across HGSOCs (SI Appendix, Fig. S1B,C, Table S1a). Notably, BORIS expression is already elevated in precursor lesions of the fallopian tube epithelium, including serous tubal intraepithelial lesions (STILs) and serous tubal intraepithelial carcinomas (STICs), suggesting that BORIS activation occurs early during HGSOC development (SI Appendix, Fig. S1D). To identify genes and pathways associated with high BORIS expression, we analyzed RNA-seq data from 11 HGSOCs with high BORIS expression and compared them with 11 tumors expressing low or undetectable levels of BORIS. Differential expression analysis identified approximately 760 genes associated with high BORIS expression, including 410 upregulated and 355 downregulated genes (Table S1b, Fig. 1C). Upregulated genes included previously reported BORIS target genes with germline-restricted expression, such as PRSS50, PRAME, and SAGE1 (25), as well as genes not previously linked to BORIS, including DPEP3, BRCA2, ACTL8, and STAG3. Genes upregulated in BORIS-high tumors were enriched for pathways related to the cell cycle, mitosis, transmembrane transport, glycoprotein hormone signaling, and meiosis, whereas genes negatively associated with BORIS expression were enriched for translational and metabolic processes (Fig. 1D, Table S1c) To extend these findings to a larger cohort and multiple ovarian cancer subtypes, we analyzed RNA-seq data from 1,202 primary different kind ovarian tumors available through cBioPortal (38). This analysis identified 238 genes that were significantly co-expressed with BORIS (Spearman correlation > 0.3, p < 0.01) (Table S1d). Of these, 47 genes overlapped with the 410 genes upregulated in BORIS-high HGSOCs (SI Appendix, Fig. S1F, Table S1e), including 15 genes associated with germ cell development and meiosis (Table S1e). Together, these findings indicate that high BORIS expression is associated with a distinct transcriptional program in ovarian cancers characterized by activation of genes involved in germline development.

The elevated BORIS expression detected in bulk RNA-seq datasets was further confirmed by analysis of published single-cell RNA-sequencing (scRNA-seq) datasets from ovarian cancer samples (39). BORIS expression was predominantly detected in transformed epithelial tumor cells, whereas little or no expression was observed in tumor-associated fibroblasts or immune cells (Fig. 1F, SI Appendix, Fig. S2A,B). The proportion of BORIS-positive cells varied considerably among HGSOC samples from different patients; however, all analyzed tumors contained at least a subset of BORIS-positive cells (SI Appendix, Fig. S2A). In some tumors, these cells formed distinct clusters, whereas in others they were more dispersed throughout the malignant cell population (SI Appendix, Fig. S2A). Consistent with the bulk RNA-seq analysis, scRNA-seq data confirmed the co-expression of several genes associated with high BORIS expression, including DPEP3, PRAME, and BRCA2 (Fig. 1F, SI Appendix, Fig. S2A,B). In addition, BORIS-positive cells expressed the canonical HGSOC markers PAX8 and WT1 (40), confirming that BORIS expression is confined to the malignant epithelial cell population (Fig. 1F, SI Appendix, Fig. S2B).

The analysis of ovarian tumor progression showed that high BORIS expression is associated with advanced-stage disease (SI Appendix, Fig. S1E). Moreover, Kaplan–Meier analysis revealed significantly poorer overall survival among ovarian cancer patients with high BORIS expression compared with those expressing low levels of BORIS (SI Appendix, Fig. S2C). Collectively, these analyses identify BORIS as a germline-specific transcription factor aberrantly reactivated early during HGSOC development, where its expression defines a distinct transcriptional program associated with poor patient outcome.

### BORIS knockout induces morphological and functional reprogramming of high-grade serous ovarian cancer cells

To investigate the functional role of BORIS in high-grade serous ovarian cancer, we generated a BORIS knockout (ko) OVCAR8 cell line, a cell model characterized by high endogenous BORIS expression (25). The CTCFL/BORIS gene produces multiple protein isoforms through the use of three promoters, alternative transcriptional start sites, and multiple translation initiation codons (41). Consequently, introducing a premature stop codon near the N-terminus would not be sufficient to eliminate all BORIS isoforms. Instead, we deleted exons 4 and 5, which encode the first four zinc fingers (ZFs) of the DNA-binding domain. Because the zinc finger domain is essential for chromatin binding and BORIS function, this strategy was designed to abolish the activity of all DNA-binding BORIS isoforms. Using the CRISPR–Cas9 system with two single-guide RNAs (sgRNAs), we introduced a ∼1,753 bp deletion encompassing ZFs 1-4 (Fig. 2A). Two single-cell clones with homozygous deletions were isolated (SI Appendix, Fig. S3A,B): clone #1 (ko1), carrying identical deletions in both alleles, and clone #2 (ko2), harboring two slightly different deletions (SI Appendix, Fig. S3A). In both clones, deletion of exons 4-5 resulted in a complete loss of full-length BORIS protein expression (Fig. 2B).

**Figure 2.**
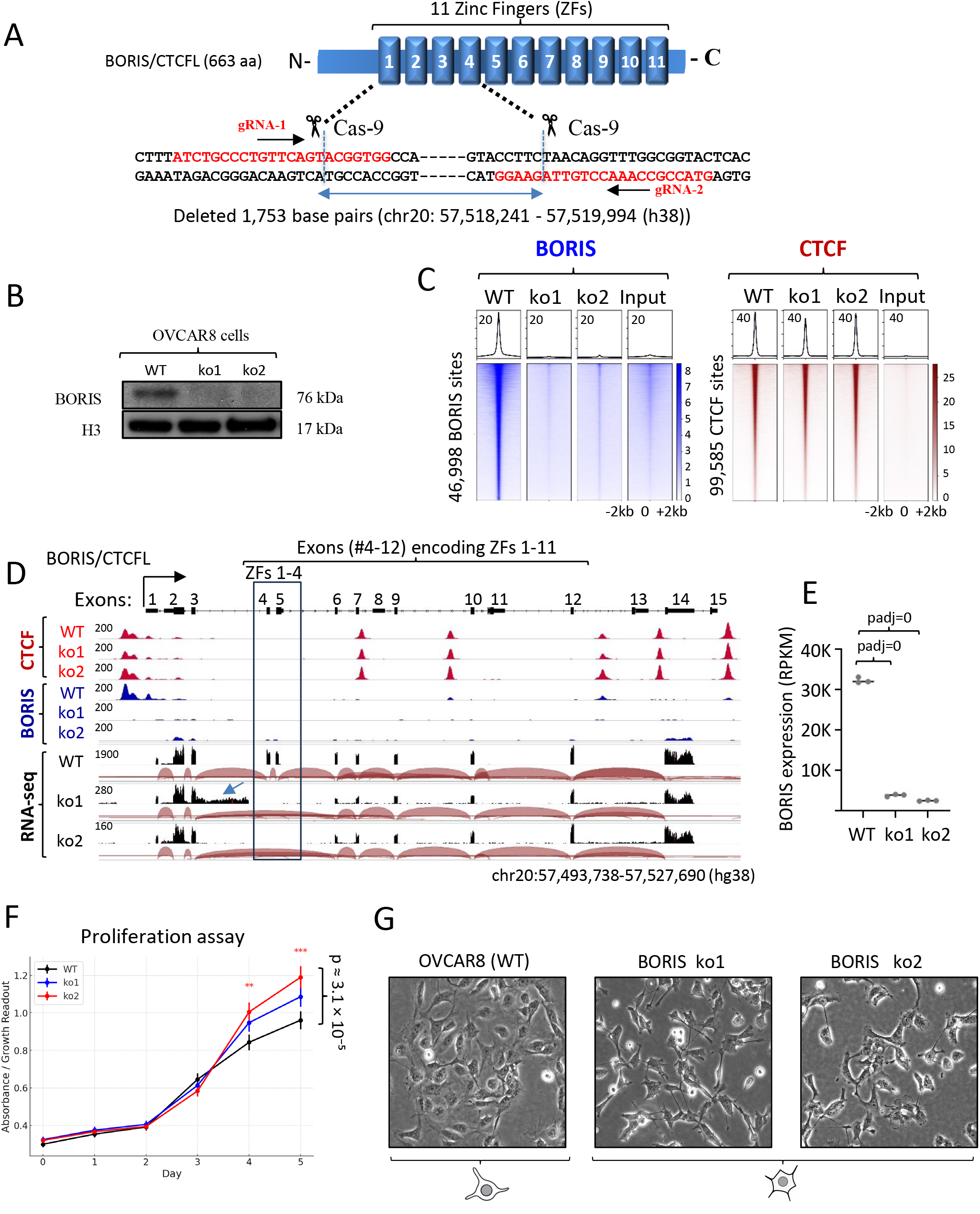
Generation and validation of BORIS knockout OVCAR8 cells. **(A)** Schematic of the CRISPR/Cas9 strategy to delete the BORIS locus in OVCAR8 cells. Two guide RNAs targeting exons encoding zinc fingers 1-4 generated a 1,753-bp deletion within the BORIS gene. **(B)** Immunoblot confirming loss of BORIS protein in two independent BORIS ko clones. **(C)** ChIP-seq showing loss of BORIS occupancy (46,998 sites) with minimal change in CTCF binding (99,585 sites). **(D)** Genome browser view of the BORIS locus showing CTCF and BORIS ChIP-seq profiles together with RNA-seq coverage in WT and BORIS ko clones. Deletion of exons 4-5, which encode zinc-fingers 1-4, abolishes BORIS expression. Blue arrow shows transcriptional readthrough from exon 3 into the downstream intron in ko1. **(E)** RNA-seq quantification showing marked reduction of BORIS transcripts in KO clones (**p < 0.001, Wald test (DESeq2)). **(F)** Proliferation assay indicating faster growth of ko clones, especially clone #2 (***p ≤ 3.1 × 10, Welch’s t-test). Data are presented as mean ± SEM. **(G)** Representative phase-contrast images of WT and BORIS ko cells illustrating morphological changes following BORIS deletion.

To evaluate how BORIS depletion affects chromatin occupancy, we performed ChIP-seq using BORIS-specific antibodies and included its paralog CTCF as a positive control (Fig. 2C), since both share similar zinc finger domains and partially overlapping genomic binding profiles (13, 25). BORIS binding was completely abolished at 46,998 sites identified in wild-type (WT) OVCAR8 cells, whereas overall CTCF occupancy remained largely unchanged (Fig. 2C). Integrative analysis of the ChIP-seq and RNA-seq datasets revealed co-occupancy of BORIS and CTCF at the BORIS promoter in OVCAR8 cells (Fig. 2D). In the ko clones, loss of BORIS binding at its own promoter was accompanied by a marked reduction in expression of the truncated BORIS transcript (Fig. 2D,E), indicating that BORIS autoregulates its own transcription.

Deletion of exons #4-5 also induced global splicing instability across the BORIS locus (SI Appendix, Fig. S3D). In ko1, transcriptional readthrough from exon #3 into the downstream intron occurred on one allele (Fig. 2D, blue arrow), generating a transcript containing a premature stop codon within the intronic sequence that does not encode a functional protein (SI Appendix, Fig. S3E). The second allele of ko1 and both alleles of ko2 predominantly spliced exon #3 directly to exon #6, with additional splicing events from exon #3 to exons #7 and #8 observed in both ko clones but not in WT cells (SI Appendix, Fig. S3D). These findings indicate that deletion of exons #4-5 disrupts normal BORIS splicing, likely compromising transcript stability, in addition to abolishing BORIS chromatin binding.

Previously, we demonstrated that complete deletion of BORIS drives differentiation of the human erythroleukemia cell line K562 toward a megakaryocytic lineage and is incompatible with K562 cell viability (25). In contrast, OVCAR8 ovarian cancer cells exhibit greater resistance to BORIS loss. Unexpectedly, both BORIS ko clones displayed increased proliferation compared with WT cells, with ko2 proliferating significantly faster than both ko1 and WT (Fig. 2F). Morphologically, both ko clones also differed from WT cells (Fig. 2G, SI Appendix, Fig. S4A). WT OVCAR8 cultures exhibited a dense, epithelial-like morphology, characterized by tightly packed, polygonal cells with prominent cell-cell junctions. In contrast, both ko clones showed increased morphological heterogeneity. ko1 cells appeared more elongated and dispersed, with reduced intercellular contacts and thinner cytoplasmic extensions, indicating a partial loss of epithelial characteristics. ko2 displayed an even more pronounced alteration, consisting of spindle-shaped, irregularly arranged cells suggestive of a mesenchymal-like phenotype (Fig. 2G).

Overall, loss of BORIS chromatin binding in OVCAR8 cells induced a morphological transition from the compact epithelial architecture typical of WT cells to a more mesenchymal and motile-like phenotype. These observations suggest that BORIS contributes to the maintenance of epithelial identity, cell adhesion, and structural organization, and that its loss promotes reprogramming of cells toward a state different from WT OVCAR8 cells.

### The knockout of BORIS in high-grade serous ovarian carcinoma results in massive transcriptional reprogramming

BORIS has been characterized as a transcriptional factor capable of epigenetically reprogramming clustered CTCF-binding sites into active promoters in both germ and cancer cells (24). Ectopic expression of BORIS in BORIS-negative somatic cells induces profound, cell context-dependent transcriptional deregulation of multiple cancer-related and germ-cell specific genes (13, 24, 42). In contrast, the consequences of BORIS loss on the transcriptional landscape of cancer cells have remained largely unexplored, with the exception of K562 leukemia cells, where BORIS downregulation was associated with terminal differentiation (25, 43).

To investigate the impact of BORIS loss in epithelial cancer, we performed transcriptomic profiling of two independent BORIS ko clones compared to WT OVCAR8 cells. Comparative analysis revealed widespread transcriptional alterations: among 21,674 genes expressed in OVCAR8 cells (baseMean > 20), up to 5,434 (25%) were differentially expressed following BORIS knockout. In ko1, 3,274 genes were upregulated and 2,106 were downregulated (adjusted p < 0.01, log FC > 1, baseMean > 20), while in ko2, 2,298 genes were upregulated and 1,220 were downregulated under the same criteria (Fig. 3A, Table S2a,b). A substantial overlap was observed between the two ko clones, with 71% of upregulated and 67% of downregulated genes changing in the same direction across both clones, although some clone-specific variations were noted as well (SI Appendix, Fig. S3E,F). Pairwise Pearson correlation analysis further confirmed that differentially expressed genes (DEGs) tended to be deregulated similarly in both ko clones (SI Appendix, Fig. S3E,F).

**Figure 3.**
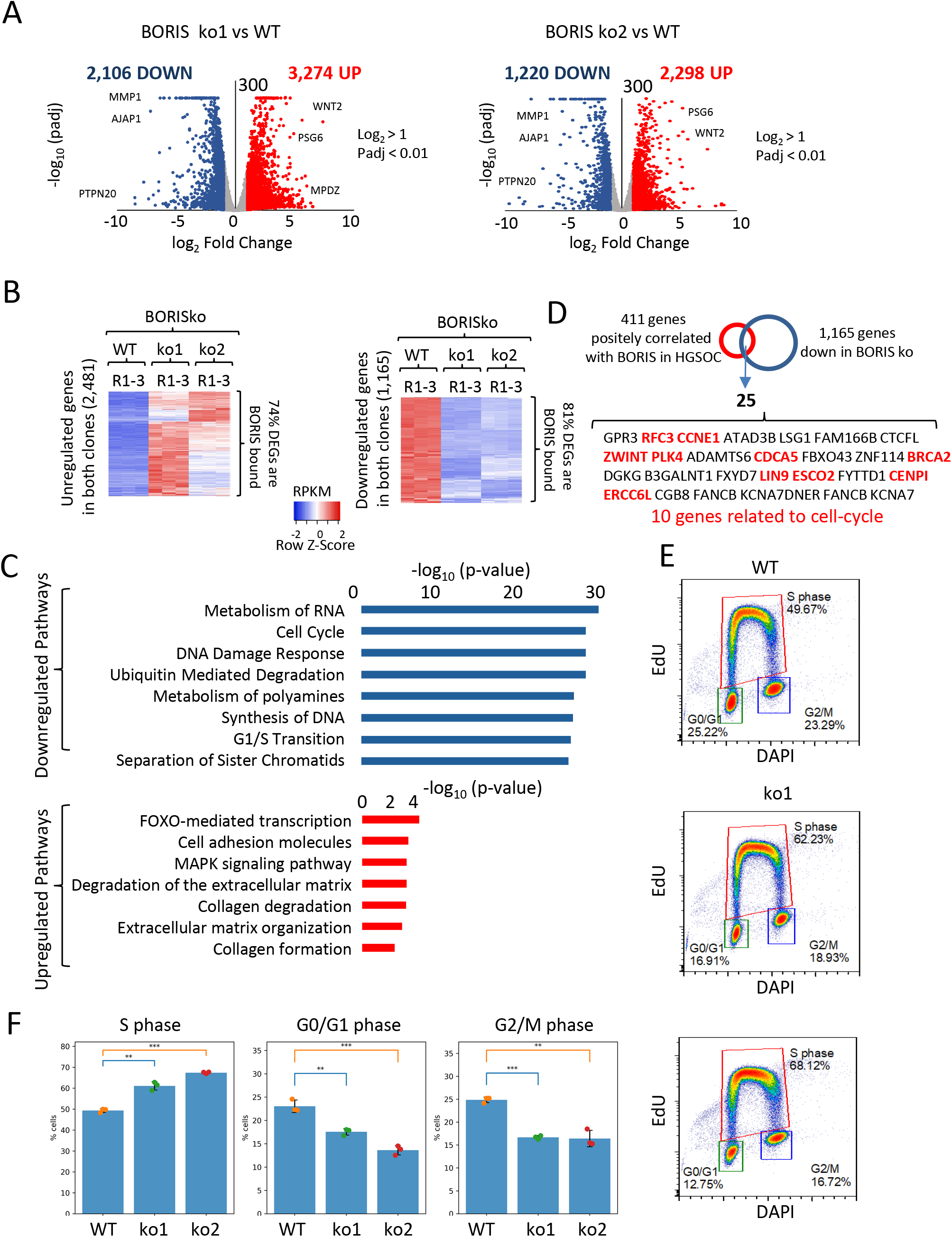
BORIS knockout induces widespread transcriptional reprogramming and cell-cycle alterations in OVCAR8 cells. **(A)** Volcano plots showing differentially expressed genes in two independent BORIS ko clones relative to WT cells. Significantly upregulated (red) and downregulated (blue) genes were identified using log fold change > 1 and adjusted P < 0.01. Selected genes are indicated. **(B)** Heatmaps of genes consistently upregulated (left) or downregulated (right) in both BORIS ko clones relative to WT. The majority of differentially expressed genes are associated with BORIS binding sites. **(C)** Pathway enrichment analysis of genes downregulated (top) and upregulated (bottom) following BORIS ko. **(D)** Overlap between genes positively correlated with BORIS expression in HGSOC and genes downregulated in BORIS ko cells identifies a common set of 25 genes, including multiple regulators of cell-cycle progression (highlighted in red). **(E)** Representative EdU incorporation and DNA content (DAPI) flow cytometry profiles of WT and BORIS ko cells showing redistribution of cells across G0/G1, S, and G2/M phases. **(F)** Quantification of the percentages of cells in S, G0/G1, and G2/M phases. Data are presented as mean ± SEM from 3 independent experiments. Statistical significance was determined using one-way ANOVA with appropriate multiple-comparison correction **p<0.001; ***p<0.0001)

To define the most robust BORIS-dependent transcriptional changes, we combined the data from both ko clones (ko1+ko2= 6 replicates) and identified 2,481 consistently upregulated and 1,165 consistently downregulated genes compared to WT OVCAR8 cells (Fig. 3B, Table S2c). Heatmap analysis demonstrated that BORIS knockout produced a more uniform pattern of gene downregulation, whereas upregulated genes exhibited greater inter-clonal variability. Importantly, the majority of both upregulated (74%) and downregulated (81%) genes overlapped significantly with BORIS-binding sites mapped in WT OVCAR8 cells (Fisher’s exact test, p < 1 × 10□³□□ and p < 6 × 10□²¹□ for up- and downregulated genes, respectively), underscoring a direct role of BORIS in transcriptional regulation of its genomic targets (Fig. 3B).

Although the number of upregulated genes was approximately twice that of downregulated genes, the downregulated pathways showed markedly higher statistical significance (top p-value < 2.47 × 10□²□), reflecting the concerted downregulation of multiple genes within specific biological processes. In contrast, upregulated pathways were less significant (top p-value < 3.98 ×10□□) and represented more heterogeneous gene sets (Fig. 3C, Table S2d). The main and significant downregulated pathways were related to RNA metabolism and the ubiquitin-proteasome pathway, including multiple proteasome subunits such as PSMAs, PSMBs, PSMCs, and PSMDs (Table S1e), all of which were severely downregulated. Moreover, their promoter analysis in respect to BORIS binding showed that these genes are all direct BORIS target (SI Appendix, Fig. S3G), suggesting that BORIS contributes to the maintenance of proteasome gene expression in ovarian cancer cells.

Among the downregulated pathways, those related to the cell cycle were the most significantly overrepresented, with 105 genes downregulated upon BORIS knockout (Fig. 3C, Table S2e). Furthermore, overlap analysis of genes positively correlated with BORIS expression in ovarian cancer (Fig. 1C, Table S1b) and genes downregulated in BORIS knockout OVCAR8 cells (Fig. 3B, Table S2c) identified 25 common genes, 10 of which are directly involved in cell-cycle progression (Fig. 3D). The connection between BORIS and cell-cycle control has been documented in several previous studies (43–45). Consistent with this, BORIS was also detected at the replication fork in OVCAR8 cells (46), implying a direct role in DNA replication. To validate these observations, we analyzed cell-cycle distribution in WT and BORIS ko cells and found that BORIS loss caused a marked accumulation of cells in the S phase, consistent with the downregulation of G1/S transition genes (Fig. 3E,F). Despite the altered cell-cycle distribution, BORIS knockout cells exhibited increased growth relative to WT control (Fig. 2F), indicating that S-phase accumulation is not associated with a proliferative arrest and may instead reflect changes in S-phase transit or replication dynamics, consistent with significant downregulation of pathways involved in cell-cycle regulation.

Among the most significantly affected pathways following BORIS knockout was extracellular matrix (ECM) organization and degradation (Table S2d). Consistent with this, many genes involved in ECM remodeling, cell adhesion, and invasion were markedly downregulated (Table S2a-c). To validate these transcriptional changes, we performed immunofluorescence staining for selected ECM-associated proteins (SI Appendix, Fig. S4A,B). Among them, LAMC2, a key component of the laminin-332 complex, was almost completely lost in both BORIS knockout clones (SI Appendix, Fig. S4, Table S2c). LAMC2 is frequently upregulated in invasive carcinomas and has been associated with enhanced cell motility, invasion, tumor progression, metastasis, and poor clinical outcome (49). Based on these observations, one might predict that BORIS ko cells would exhibit reduced tumorigenic potential. Surprisingly, however, BORIS ko cells proliferated significantly faster than WT cells (Fig. 2F), in contrast to previous studies implicating BORIS in tumor progression (24, 30, 45). This unexpected phenotype suggests that the consequences of BORIS loss extend beyond the downregulation of individual pro-tumorigenic genes. For example, AJAP1 (Adherens Junctions Associated Protein 1), a protein involved in cell adhesion, cell-ECM interactions, and invadopodia regulation (50), was also strongly downregulated in both BORIS ko clones (SI Appendix, Fig. S4, Table S2c). Together, these findings indicate that BORIS loss broadly remodels ECM and adhesion pathways, producing complex cellular phenotypes that cannot be explained by changes in a single downstream target.

To further assess how BORIS loss influences the malignant properties of OVCAR8 cells, we performed several functional assays. In the foci formation assay, WT and BORIS ko cells formed a similar number of colonies of comparable size (SI Appendix, Fig. S5A,B). However, WT colonies exhibited more intense crystal violet staining (SI Appendix, Fig. S5C), consistent with a higher cell density within individual colonies. Because WT cells proliferated more slowly than BORIS knockout cells (Fig. 2F), this observation may reflect enhanced cell-cell adhesion or reduced contact inhibition in WT cells. In contrast, BORIS ko2 cells closed the wound significantly faster than WT OVCAR8 cells in the scratch assay, indicating increased migratory capacity (SI Appendix, Fig. S5D,E).

We next examined the response of BORIS ko cells to cisplatin and carboplatin (SI Appendix, Fig. S5F,G). Although the BORIS ko1 clone exhibited significantly greater resistance to platinum treatment than WT cells, no comparable effect was observed in the BORIS ko2 clone, suggesting that increased chemoresistance reflects clonal variation rather than a direct consequence of BORIS loss. Overall, despite widespread transcriptional deregulation of cancer-associated genes following BORIS depletion, BORIS knockout did not reduce the malignant properties of OVCAR8 cells and, in some assays, was associated with enhanced migratory behavior.

### BORIS loss remodels the epigenetic landscape and destabilizes CTCF-cohesin occupancy

To determine how BORIS loss drives widespread transcriptional reprogramming, we examined changes in chromatin accessibility, active histone modifications, and CTCF/cohesin occupancy. Although CTCF and BORIS recognize the same DNA motif in vitro, their genomic binding profiles in OVCAR8 cells are only partially overlapping, with most CTCF-binding sites lacking BORIS occupancy (SI Appendix, Fig. S6A,B). Consistent with our previous observations (25), genomic regions co-bound by CTCF and BORIS, as well as BORIS-only sites, are enriched for clustered CTCF-binding sites (2×CTSes) and active chromatin marks, including H2A.Z, H3K4me3, and H3K27ac (SI Appendix, Fig. S6C-E). We therefore compared the genome-wide distribution of these active histone marks together with CTCF, BORIS, and cohesin (RAD21 and SMC3) occupancy in WT and BORIS ko cells. We first examined representative genes that were consistently deregulated in both ko clones before extending our analysis to genome-wide chromatin changes.

BORIS has been described as a transcriptional activator of testis-specific genes when ectopically expressed in BORIS-negative somatic cells, supporting its role as a regulator of germline transcriptional programs (24, 42). We therefore asked whether BORIS loss in cancer cells leads to the silencing of aberrantly activated testis-specific genes. Indeed, one of the most striking examples was PTPN20 (protein tyrosine phosphatase, non-receptor type 20), a predominantly testis-expressed gene that became completely silenced upon BORIS loss (Table S2c, Fig. 4A). This transcriptional shutdown coincided with the loss of BORIS binding at the PTPN20 promoter, accompanied by a simultaneous loss of CTCF and cohesin occupancy and depletion of active chromatin marks (H2A.Z, H3K4me3, H3K27ac) (Fig. 4A, red arrows). Thus, BORIS binding appears essential for maintaining the active chromatin configuration and transcriptional outcome of the PTPN20 promoter.

**Figure 4.**
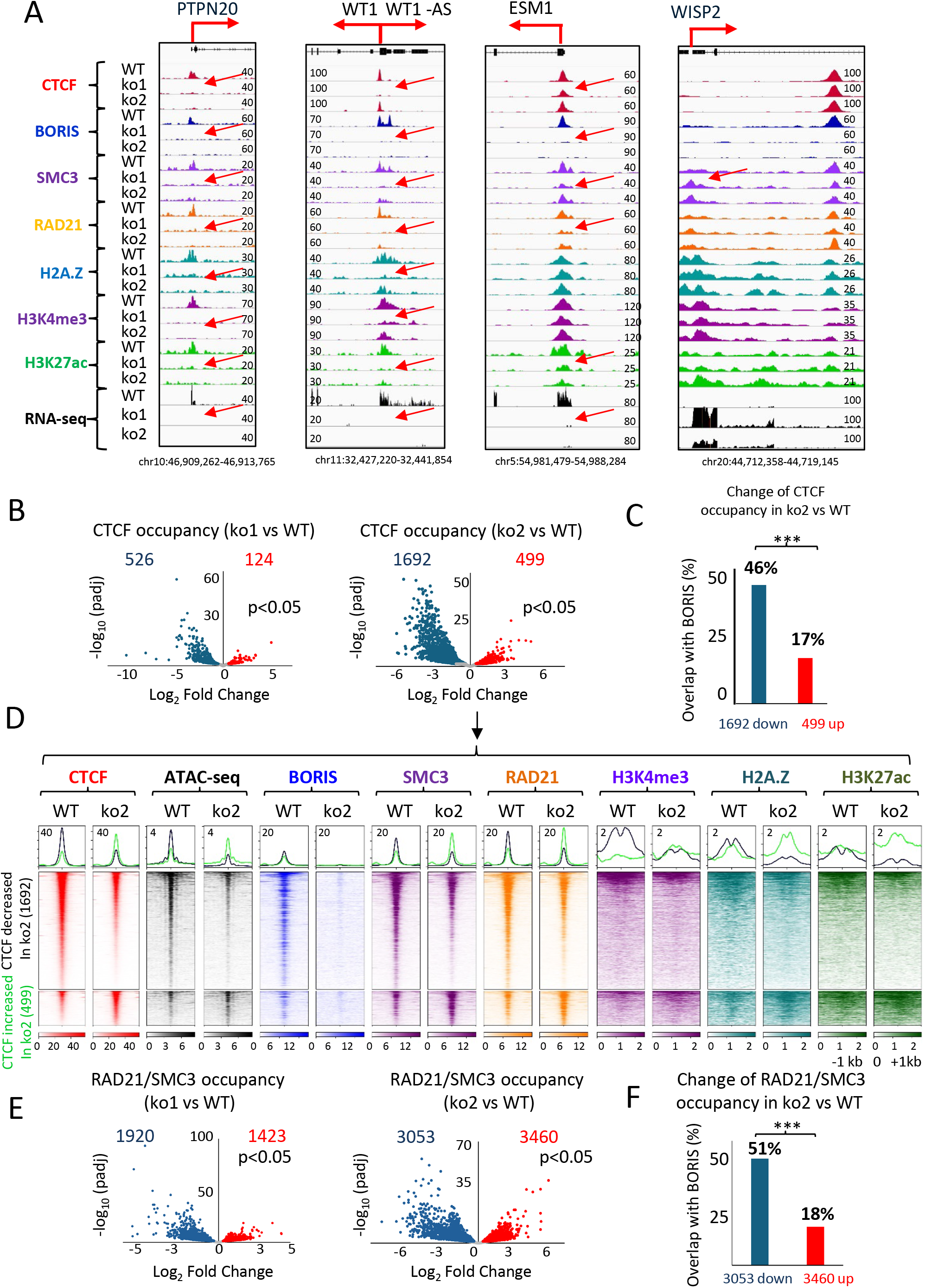
BORIS knockout remodels chromatin occupancy and epigenetic landscapes in OVCAR8 cells. **(A)** Representative genome browser views showing CTCF, BORIS, SMC3, RAD21, H2A.Z, H3K4me3, and H3K27ac ChIP-seq profiles together with RNA-seq coverage at selected loci in WT cells and two independent BORIS ko clones. Red arrows indicate BORIS-bound regions that exhibit altered chromatin occupancy and gene expression changes following BORIS ko. **(B)** Volcano plots showing genomic regions with significantly decreased (blue) or increased (red) CTCF occupancy in BORIS ko clones relative to WT. **(C)** Fraction of regions (%) with altered CTCF occupancy that overlap BORIS-binding sites. **(D**) Heatmaps and average ChIP-seq signal profiles of CTCF, ATAC-seq, BORIS, SMC3, RAD21, H3K4me3, H2A.Z, and H3K27ac centered on genomic regions showing decreased (top) or increased (bottom) CTCF occupancy in ko2 cells relative to WT. Signals are displayed within ±1 kb of the summit of CTCF ChIP-seq peak. **(E)** Volcano plots showing genomic regions with significantly decreased (blue) or increased (red) RAD21/SMC3 occupancy in BORIS ko clones relative to WT. **(F)** Fraction of regions (%) with altered RAD21/SMC3 occupancy that overlap BORIS-binding sites in WT cells. *** Statistical significance was assessed using the chi-square test (p< 0.001).

Another example of a direct BORIS target is WT1 (Wilms Tumor 1), one of the most specific and reliable biomarkers of high-grade serous ovarian carcinoma (51). WT1 is implicated in cell proliferation, epithelial-mesenchymal transition (EMT), metastasis, stemness, and chemoresistance in ovarian cancer (52). Analysis of primary ovarian tumors revealed a strong positive correlation between BORIS and WT1 expression in HGSOC (Fig. 1F, SI Appendix, Fig. S2B, Table S2c), suggesting that WT1 expression is associated with BORIS expression in malignant contexts. Integration of multiple next-generation sequencing (NGS) datasets demonstrated that loss of BORIS occupancy at the WT1 promoter resulted in complete transcriptional silencing of WT1 (Fig. 4A). This was accompanied by complete loss of CTCF binding in ko1 cells, whereas CTCF occupancy was partially retained in ko2 cells. Despite these clone-specific differences, both BORIS ko clones exhibited loss of cohesin occupancy, depletion of active histone marks, and complete silencing of WT1 (Fig. 4A, red arrows). BORIS depletion also abolished expression of WT1-AS, the antisense transcript that is normally co-expressed with WT1 (53). Together, these findings identify WT1 as a direct BORIS-regulated gene whose expression depends on BORIS-mediated maintenance of an active chromatin state in some cells.

A third example is ESM1 (Endothelial Cell-Specific Molecule 1), a secreted proteoglycan implicated in angiogenesis and tumor progression and increasingly recognized as a pro-tumorigenic and prognostic biomarker across multiple cancers, including ovarian cancer (54). ESM1 promoter belongs to the class of CTCF/BORIS co-bound sites and represents a direct BORIS target (Fig. 4A). While CTCF occupancy remained at the ESM1 promoter following BORIS loss, cohesin binding decreased markedly in both ko clones (Fig. 4A, red arrows), accompanied by depletion of H3K27ac. Taken together, these examples illustrate that BORIS loss frequently leads to destabilization of local chromatin architecture, loss of CTCF and cohesin occupancy, and depletion of active epigenetic marks at its target loci, resulting in transcriptional silencing of key genes.

Conversely, WISP2 (WNT1-inducible signaling pathway protein 2), a member of the CCN family involved in cell growth, migration, differentiation, and extracellular matrix remodeling (55), was among the most strongly upregulated genes following BORIS ko. Consistent with previous reports showing that ectopic BORIS expression represses WISP2 in MCF7 breast cancer cells (25), WISP2 expression negatively correlated with BORIS levels in HGSOC tumors (Fig. 1C, Table S1a) and was markedly increased in both BORIS ko clones. Although neither CTCF nor BORIS binds directly to the WISP2 promoter, BORIS occupies a downstream regulatory region, while cohesin occupancy increased at the WISP2 promoter following BORIS depletion (Fig. 4A, red arrow). These findings suggest that BORIS regulates WISP2 through distal chromatin interactions rather than direct promoter binding.

To determine whether these gene-specific changes extend to the genome-wide level, we examined the effects of BORIS depletion on CTCF occupancy. Hundreds of CTCF-binding sites were either lost or gained in both BORIS ko clones relative to WT cells (Fig. 4B). Notably, the loss of CTCF occupancy was significantly enriched at BORIS-bound regions, with 46% of CTCF sites that lost occupancy overlapping BORIS-binding sites in WT OVCAR8 cells (Fig. 4C). These findings suggest that BORIS contributes to maintaining CTCF occupancy at a subset of genomic loci. In contrast, newly acquired CTCF-binding sites showed much less (17%) overlap with BORIS-bound regions (Fig. 4C), indicating that BORIS depletion redistributes, rather than globally reduces, CTCF occupancy.

To integrate this data with chromatin status, we performed a pairwise heatmap analysis comparing sites with either lost or gained CTCF occupancy after BORIS ko (Fig. 4D, SI Appendix, Fig. S7A). As expected, loss or gain of CTCF binding was accompanied by corresponding changes in chromatin accessibility (ATAC-seq): regions losing CTCF binding showed reduced accessibility, whereas regions gaining CTCF binding became more open. Consistent with these patterns, the occupancy of cohesin subunits (SMC3, RAD21) and the levels of active histone modifications (H3K4me3, H2A.Z, H3K27ac) changed in the same direction as CTCF binding (Fig. 4D, SI Appendix, Fig. S7A).

Interestingly, although BORIS is not able to interact directly with cohesin (12), cohesin occupancy appeared particularly sensitive to BORIS loss in the individual gene examples (Fig. 4A). We therefore examined cohesin binding genome-wide in both ko clones. Notably, at least twice as many cohesin-binding sites changed occupancy following BORIS ko compared with CTCF-binding sites (Fig. 4E). Similar to CTCF, loss of cohesin binding was significantly enriched at BORIS-bound regions (Fig. 4F), suggesting that BORIS contributes indirectly to the stabilization of CTCF-bound chromatin domains. In the absence of BORIS, destabilization of CTCF occupancy weakens cohesin binding, potentially compromising the strength and stability of chromatin loop anchors.

We previously showed that BORIS can recruit the SRCAP complex, facilitating the replacement of canonical histone H2A with its variant H2A.Z and thereby promoting local chromatin accessibility and activation of alternative promoters via CTCF binding sites (24). Consistent with this mechanism, BORIS ko in OVCAR8 cells led to the most pronounced reduction in H2A.Z enrichment specifically at BORIS-bound sites (BORIS-only and CTCF/BORIS co-bound regions) compared with other active histone marks (SI Appendix, Fig. S7B). Taken together, these findings reveal that BORIS is essential for maintaining the structural and epigenetic integrity of CTCF-associated chromatin domains. Its loss compromises CTCF and cohesin binding, alters chromatin accessibility and active histone modifications, and consequently drives large-scale transcriptional reprogramming in OVCAR8 cells.

### BORIS loss induces coordinated transcriptional changes within genomic gene clusters

A more detailed analysis of the RNA-seq data revealed that many differentially expressed genes were organized into genomic clusters that were coordinately deregulated in both BORIS ko clones, suggesting non-random regulation at the level of chromatin domains. One striking example was the pregnancy-specific glycoprotein (PSG) locus on chromosome 19q13.2, where all functional PSG genes (PSG1-PSG9 and PSG11) together with 33 neighboring genes were upregulated (Fig. 5A,B). Because PSG genes are normally restricted to placental tissues (56), their coordinated activation suggests locus-wide derepression, likely reflecting changes in local chromatin organization.

**Figure 5.**
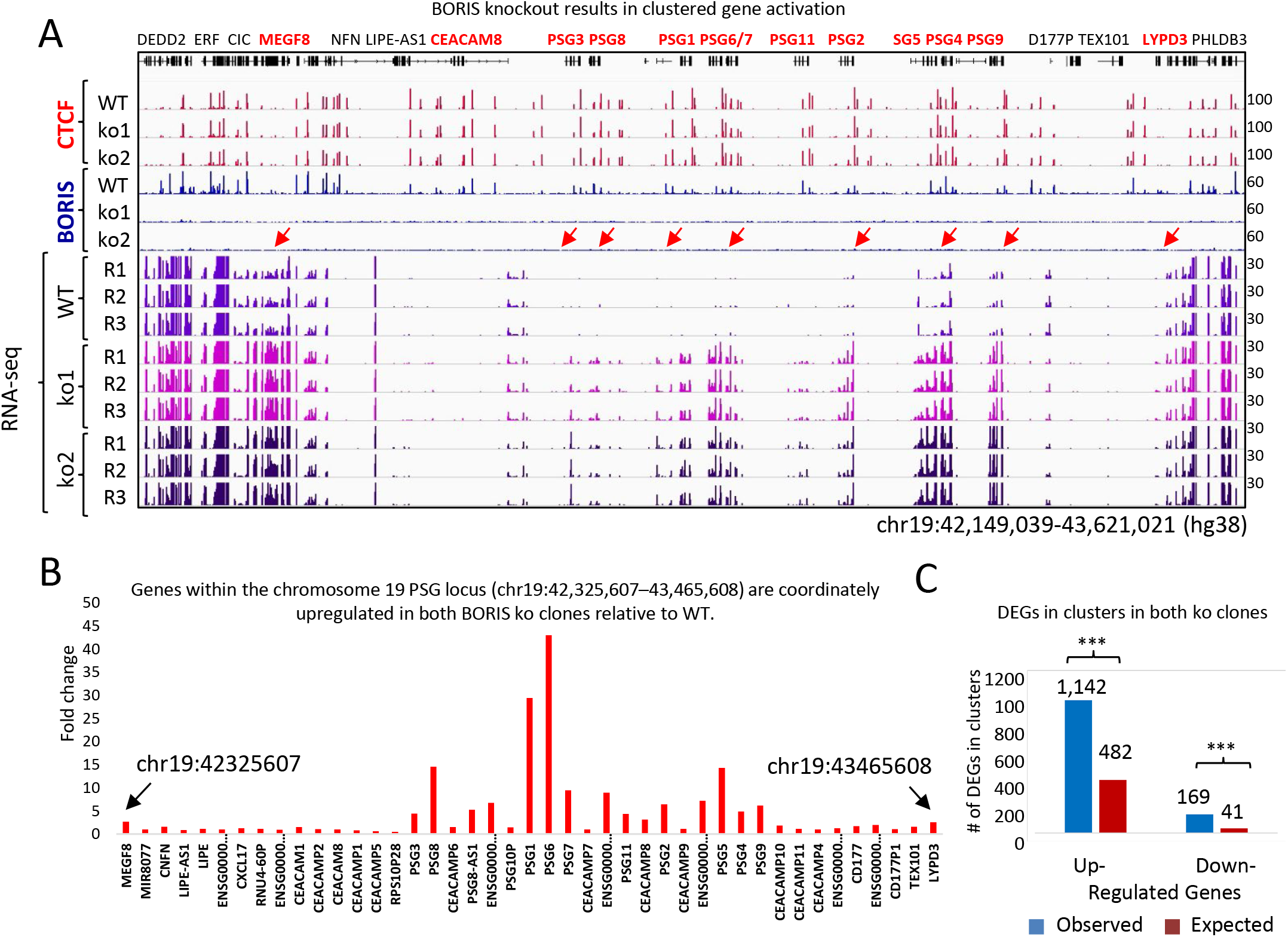
BORIS knockout induces coordinated activation of genes in clusters. **(A)** Genome browser view of the chromosome 19 locus (chr19:42,149,039-43,621,021; hg38) showing CTCF and BORIS ChIP-seq profiles together with RNA-seq coverage in WT cells and two BORIS ko clones. Red arrows indicate genes within the pregnancy-specific glycoprotein (PSG) cluster that are coordinately upregulated following BORIS deletion. **(B)** Log fold changes in expression of genes across the same genomic interval in BORIS ko clones relative to WT, demonstrating coordinated upregulation of the PSG gene cluster. **(C)** Observed and expected numbers of differentially expressed genes (DEGs) located within genomic gene clusters among genes consistently upregulated or downregulated in both BORIS ko clones. Statistical significance was assessed using the chi-square test (*** p<0.004).

To determine whether differential gene expression following BORIS knockout was non-randomly distributed across the genome, we performed a positional clustering analysis using a 100-kb proximity threshold. Using stringent differential expression criteria (log FC > 1, P < 0.001), we analyzed upregulated and downregulated genes separately. We identified 339 clusters containing 1,142 upregulated DEGs and 48 clusters containing 169 downregulated DEGs (Table S3a). Permutation analysis (250 iterations) demonstrated that the number of clustered genes, the number of clusters, and the maximum cluster size were all significantly greater than expected by chance (Fig. 5C). Consistently, clustered DEGs were separated by significantly shorter genomic distances than predicted by the null distribution. Twenty-one genomic intervals exhibited significant clustering, including hotspots encompassing the HLA, protocadherin, PSG, and kallikrein gene families (Table S3a). Together, these findings indicate that BORIS loss induces coordinated transcriptional changes within localized genomic domains, consistent with chromatin-level regulation.

The deregulation of clustered genes was also observed in a clone-specific manner (Table S3b,c). For example, a set of 15 genes within the chromosome 19p12 locus (chr19: 22,174,764-22,686,732) was highly and significantly upregulated in ko2, but not in ko1, relative to WT cells (SI Appendix, Fig. S8A,B; Table S3b,c). Notably, similar to the PSG gene family, many of the genes in this locus belong to the zinc finger protein family (ZNF676, ZNF729, ZNF98, ZNF209P, ZNF492, ZNF849P, ZNF99, ZNF723, ZNF730) (Table S3c). Although this genomic region is not well characterized in the literature, several of these ZNF genes, such as ZNF676, are primate-specific and have been implicated in repressing subsets of endogenous retroviral long terminal repeat (LTR) elements, particularly the LTR12C family, in the germline and early embryo (57). Their coordinated upregulation is also noteworthy because several members of this cluster (VN1R85P, RAD54L2P1, ZNF676, ZNF729, ZNF98, ZNF492) are highly expressed in testis and may normally be regulated by BORIS during spermatogenesis. Thus, given that BORIS is a paralog of the 3D chromatin organizer CTCF, the appearance of clustered gene deregulation suggests that BORIS loss may disrupt higher-order chromatin architecture.

### BORIS depletion strengthens A/B chromatin compartmentalization

To determine whether BORIS depletion alters higher-order chromatin organization, we performed Hi-C analysis of WT and OVCAR8 ko2 cells. Hi-C contact maps revealed multiple chromosomal fusions and rearrangements consistent with the known karyotype of OVCAR8 cells (SI Appendix, Fig.S9A,B). Analysis of distance-dependent contact probability showed highly similar profiles across biological replicates and resolutions, with near-complete overlap between WT and ko cells across the entire range of genomic distances examined (Fig. 6A, SI Appendix, Fig. S9C). No consistent shift in the decay curve was observed, indicating that global chromatin compaction and large-scale genome folding remained largely unchanged following BORIS ko (Fig. 6A). Consistent with this observation, the overall numbers of cis and trans interactions and long-range cis contacts were similar between WT and ko cells (SI Appendix, Fig. S9D,E).

**Figure 6.**
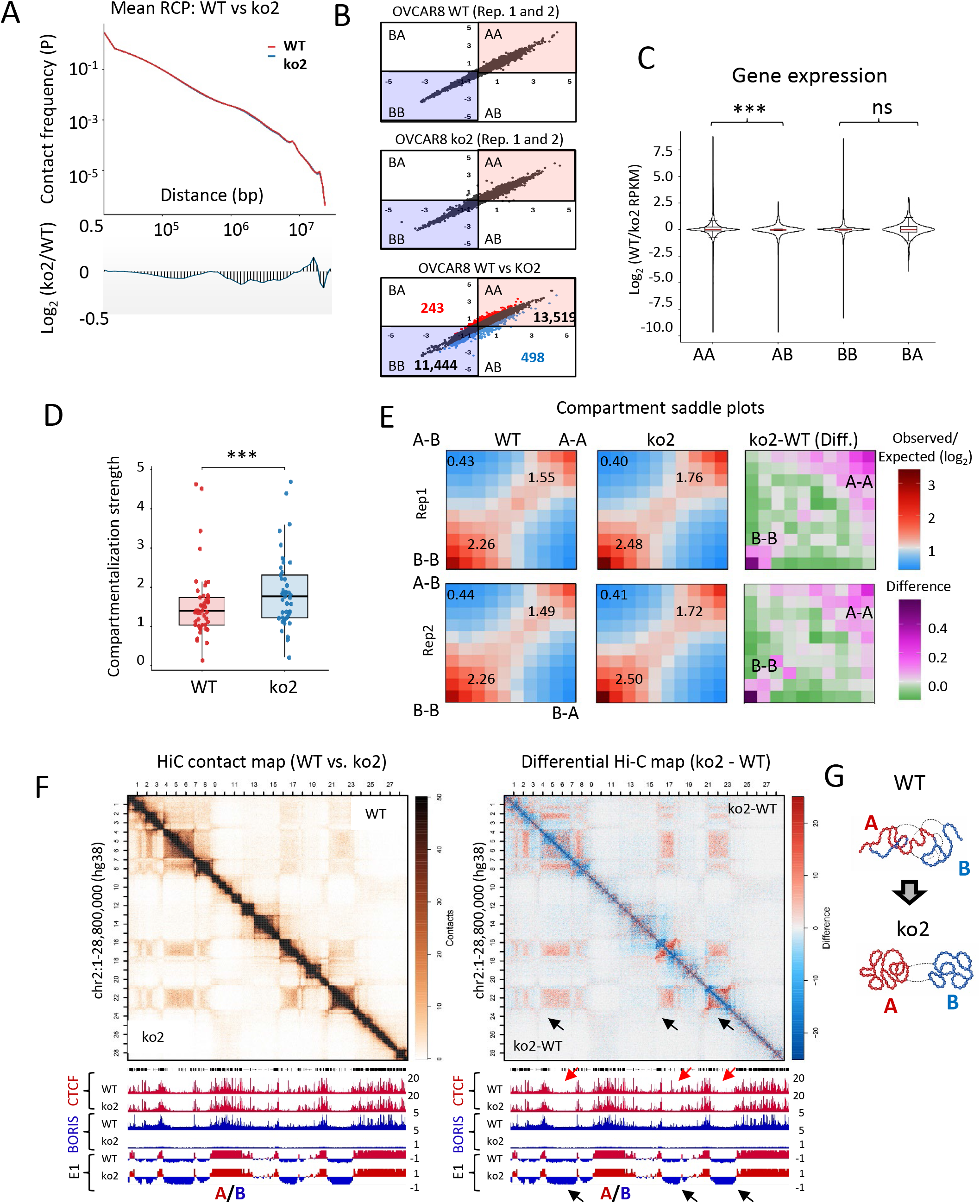
BORIS knockout increases chromatin compartmentalization in OVCAR8 cells. **(A)** Mean contact probability (P(s)) curves for WT and BORIS ko2 Hi-C datasets (top). The lower panel shows the log ratio of contact probability (ko2/WT) as a function of genomic distance. **(B)** Scatter plot of E1 compartment scores comparing WT and ko2 cells. Red and blue numbers indicate the numbers of genomic bins with significantly increased or decreased E1 scores, respectively (p < 0.05), following BORIS ko, whereas black numbers denote bins that did not exhibit significant changes in compartment identity. **(C)** Violin plots overlaid with boxplots showing the distribution of gene expression changes (log RPKM, WT/ko2) for genes located in AA, AB, BB, and BA compartment classes. Statistical significance was assessed using the Welch’s t-test (*** (P = 1.6 × 10); ns (P = 0.486)). **(D)** Quantification of chromatin compartment strength in WT and ko2 cells. *** - Statistical significance was assessed using paired Wilcoxon test (p = 2.29e-09) and paired t-test (p = 7.36e-09). **(E)** Saddle plots for two biological replicates of WT and ko2 cells together with difference maps (ko2-WT), illustrating increased segregation of A-A and B-B interactions following BORIS ko. **(F)** Representative HiC contact maps (left) and differential interaction map (center; ko2-WT) showing increased compartmentalization following BORIS ko. Black and red arrows indicate representative regions showing strengthening of the B compartment and concomitant loss of CTCF occupancy, respectively. **(G)** A schematic summarizes the increased segregation of A and B chromatin compartments in BORIS ko cells.

Despite the absence of global changes in chromatin folding, compartment analysis at 100-kb resolution revealed substantial reorganization of A/B compartment structure following BORIS ko. A total of 498 genomic bins displayed a significant shift toward the B compartment (A→B direction), whereas 243 shifted toward the A compartment (B→A direction). (Fig. 6B). Genes located within AB compartments were significantly downregulated compared with genes residing in stable AA compartments (Fig. 6C). In contrast, genes within BA compartments showed a trend toward increased expression relative to genes in stable BB compartments, although this difference did not reach statistical significance (Fig. 6C).

Beyond compartment switching, BORIS depletion produced a definite increase in compartmentalization strength. Analysis of the first eigenvector (E1) values revealed significantly stronger compartment segregation in ko cells compared with WT cells (Fig. 6D). Consistently, saddle plot analysis demonstrated enhanced A-A and B-B interactions together with reduced A-B mixing following BORIS ko, indicating stronger spatial segregation of active and inactive chromatin compartments within the nucleus (Fig. 6E). This effect was particularly pronounced for B compartments, which exhibited substantially stronger B–B interactions in ko2 cells than in WT cells (Fig. 6E,F). Representative genomic regions illustrating increased B–B compartment interactions are shown in Fig. 6F, Appendix, Fig. S10A. Notably, the strengthening of interactions among B compartments in ko2 cells also displayed reduced CTCF occupancy at several sites (red arrows, Fig. 6F, Appendix, Fig. S10A), suggesting a relationship between BORIS loss, altered CTCF binding, and enhanced compartment segregation. Increased compartmentalization was also associated with coordinated transcriptional changes across extended genomic regions. For example, activation of the PSG gene cluster (Fig. 5) correlated with a significant increase in A-compartment strength across the entire locus in BORIS ko cells (Appendix, Fig. S9F). Together, these findings indicate that BORIS depletion reinforces compartmental segregation, resulting in stronger separation of active and inactive chromatin domains, as summarized schematically in Fig. 6G.

### BORIS depletion increases TAD insulation

The next level of chromatin organization after A/B compartments is the topologically associating domain (TAD). To determine whether BORIS depletion affects TAD organization, we first performed quantitative TAD boundary analysis using insulation score-based boundary strength measurements. Differential analysis identified 396 significantly remodeled TAD boundaries in BORIS ko2 cells compared to WT cells, including 238 strengthened (gained) and 158 weakened boundaries (lost), whereas the vast majority of boundaries (7,042) remained unchanged (shared) (Fig. 7A). These findings indicate that BORIS depletion does not globally reorganize TAD architecture.

**Figure 7.**
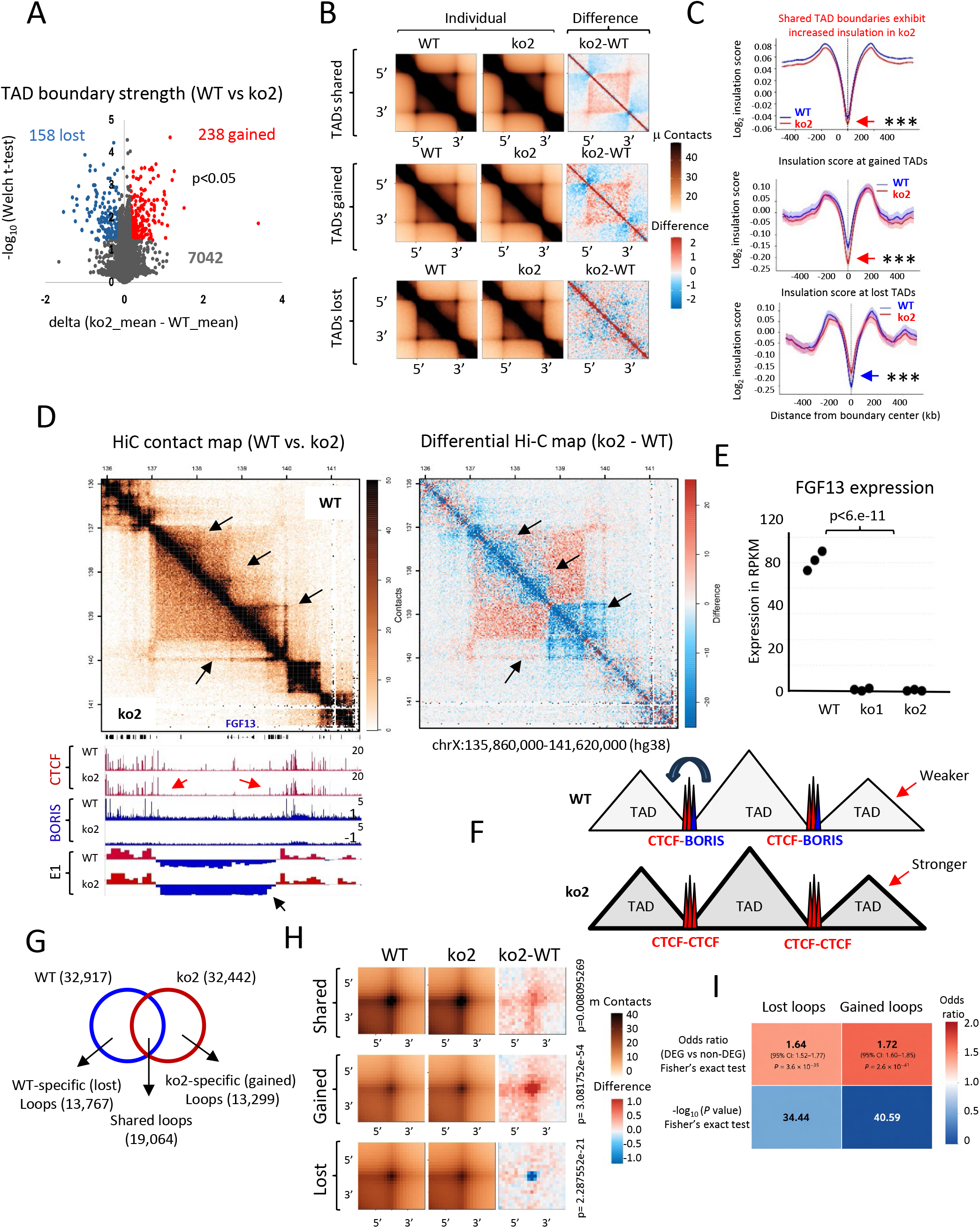
BORIS regulates TAD boundary strength and chromatin loop organization. **(A)** Volcano plot showing changes in TAD boundary strength following BORIS ko relative to WT. Significantly strengthened (red) and weakened (blue) TAD boundaries were identified using Welch’s t-test (p < 0.05). **(B)** Aggregate TAD analyses comparing shared, gained, and lost TAD boundaries in WT and ko2 cells. Difference maps (ko2-WT) highlight changes in local chromatin interactions. **(C)** Average insulation score profiles centered on shared, gained, and lost TAD boundaries. Shared TAD boundaries exhibit increased insulation following BORIS ko, whereas gained and lost boundaries display corresponding changes in insulation strength. Shaded areas represent the SEM. Statistical significance was assessed using the paired Wilcoxon test (p< 1×10 ^5^). **(D)** Representative HiC contact maps (left) and differential interaction map (center; ko2-WT) illustrating changes in TAD organization following BORIS ko. Corresponding CTCF and BORIS ChIP-seq profiles and compartment score (E1) tracks are shown below. Black arrows indicate representative changes in compartmentalization and TAD boundary strength, whereas red arrows indicate regions exhibiting reduced CTCF occupancy in ko2 cells. **(E)** RNA-seq quantification of FGF13 expression in WT and BORIS ko clones. **(F)** Model illustrating BORIS-dependent regulation of TAD boundaries. In WT cells, BORIS occupancy weakens CTCF-mediated boundary insulation, whereas following BORIS ko, CTCF occupancy strengthens TAD boundaries. **(G)** Overlap of chromatin loops identified in WT and ko2 cells, showing shared, lost, and gained loops following BORIS ko. **(H)** Aggregate peak analysis of shared, gained, and lost chromatin loops in WT and ko2 cells. Difference maps (ko2–WT) illustrate changes in loop strength. P values, determined by the Wilcoxon test, are shown on the left. **(I)** Enrichment analysis showing that differentially expressed genes (DEGs) are significantly associated with both lost and gained chromatin loops. Odds ratios, 95% confidence intervals, and Fisher’s exact test P values are indicated.

Approximately 28% of BORIS binding sites and 26% of CTCF binding sites overlapped TAD boundaries in OVCAR8 cells. Conversely, 74% of TAD boundaries were occupied by BORIS and 97% by CTCF, indicating that BORIS is frequently bound at TAD boundaries together with CTCF. Because BORIS, unlike CTCF, is not able to block cohesin-mediated loop extrusion (12), these observations suggest that BORIS occupancy may have an impact on TAD insulation. To further investigate this possibility, we examined the occupancy of CTCF, RAD21, and BORIS at gained, lost, and shared TAD boundaries (SI Appendix, Fig. S10B). BORIS was enriched together with CTCF and RAD21 at all classes of TAD boundaries. Following BORIS depletion, CTCF and RAD21 occupancy were reduced specifically at lost TAD boundaries, whereas occupancy at shared and gained boundaries remained largely unchanged (SI Appendix, Fig. S10B). These results suggest that BORIS contributes to the stability of a subset of CTCF/cohesin-bound TAD boundaries, as was shown in Fig. 4.

To determine how BORIS loss affects TAD insulation, we generated aggregate TAD pileups for boundaries shared between WT and ko2 cells, as well as TADs gained or lost following BORIS depletion. Overall, TAD architecture was largely preserved; however, aggregate pileups consistently revealed stronger intra-TAD interactions in ko2 cells than in WT cells (Fig. 7B,C). Difference maps (ko2/WT) of shared TADs showed increased contact frequencies within TADs accompanied by reduced interactions across TAD boundaries, indicating that BORIS depletion enhances TAD insulation even at boundaries that remain unchanged between WT and ko2 cells (Fig. 7B, upper panel). Consistent with the pileup analysis, insulation score profiles at shared TAD boundaries showed a highly significant increase in boundary insulation following BORIS ko (paired Wilcoxon test, P < 1 × 10 ³). As expected, boundaries classified as lost or gained displayed decreased or increased insulation, respectively (Fig. 7B,C).

Comparison of TAD organization with compartment strength demonstrated a close relationship between these two levels of genome organization. Regions showing increased TAD insulation frequently also exhibited stronger compartment segregation and corresponding changes in gene expression (Fig. 6F, 7D,E, SI Appendix, Fig. S10A). One representative example is the FGF13 locus (Fig. 7D,E). FGF13 (Fibroblast Growth Factor 13), which plays important roles in neuronal development and electrical signaling (58), is expressed in WT OVCAR8 cells but it is silenced following BORIS knockout in both ko clones (Fig. 7E). FGF13 silencing in ko2 cells was accompanied by strengthening the surrounding B compartment, loss of CTCF occupancy within the B compartment (red arrows, Fig. 7D), fusion of adjacent subTADs into a single larger insulated TAD, and a marked increase in TAD insulation (Fig. 7D, right panel). A similar pattern is shown at additional loci (Fig. 6F, SI Appendix, Fig. S10A), where enhanced B-compartment segregation was associated with loss of internal CTCF binding, disappearance of subTAD organization, and strengthening of TAD insulation.

Across 74% loci, BORIS co-occupied TAD boundaries together with CTCF in WT cells, whereas only CTCF remained bound following BORIS depletion in ko2 cells. These observations suggest that BORIS normally attenuates TAD boundary insulation, and that its loss results in stronger, more insulated TAD boundaries (schematic summary presentation in 7F). This model is consistent with the inability of BORIS to block cohesin-mediated loop extrusion and provides a mechanistic explanation for the increased chromatin insulation observed following BORIS depletion.

### BORIS depletion increases loop strength

At a finer scale of genome organization (5-25 kb resolution), the total number of chromatin loops identified in WT and ko2 cells was similar (approximately 32,000 loops in each condition). However, fewer than 60% of loops were shared between WT and ko2 cells, indicating substantial loop rewiring following BORIS depletion (Fig. 7G). Similar to the TAD analysis, Aggregate Peak Analysis (APA) revealed that both shared loops and ko2-specific loops exhibited significantly stronger interaction intensities in BORIS ko cells than in WT cells (Fig. 7H). These findings indicate that although BORIS depletion does not alter the overall number of chromatin loops, it strengthens both shared and ko-specific chromatin loops while promoting extensive rewiring of chromatin looping interactions.

To determine whether loop stability was associated with CTCF and BORIS occupancy, loops were classified according to the highest-priority binding event detected at either anchor. Loop categories were assigned hierarchically based on the highest-priority CTCF/BORIS annotation detected at either loop anchor, followed by CTCF-only occupancy, BORIS-only occupancy, and the absence of both factors. Shared loops were enriched for anchors containing CTCF/BORIS co-bound sites (59.6%), whereas WT-specific and ko-specific loops exhibited substantially lower frequencies (50.8% and 49.5%, respectively) (SI Appendix, Fig. S10C). This suggests that loops associated with CTCF/BORIS are generally more stable. Cell-specific loops displayed a marked increase in anchors lacking detectable CTCF or BORIS occupancy, rising from 10.0% among shared loops to 16.7–17.8% among condition-specific loops (SI Appendix, Fig. S10C). CTCF-only anchor occupancy was largely preserved across loop classes, implying that the major distinction is not loss of CTCF itself (SI Appendix, Fig. S10C). BORIS-only anchors represented less than 1% of loops in all categories and more likely a result of random overlapping (SI Appendix, Fig. S10C), supporting the model that BORIS acts primarily at CTCF-associated sites, not through an independent BORIS-only loop network. We further asked whether loop stability was associated with the occupancy status of both anchors. Shared loops were enriched for loops anchored by CTCF/BORIS-associated sites at both ends, whereas WT-specific and KO-specific loops showed reduced frequencies of such configurations (SI Appendix, Fig. S10D). These findings suggest that CTCF/BORIS-associated loops preferentially belong to the conserved loop repertoire and may constitute a stable architectural scaffold within the genome.

To assess the relationship between transcriptional changes and 3D genome remodeling, we examined the overlap between differentially expressed genes (DEGs), non-differentially expressed genes (non-DEGs), and condition-specific chromatin loops (Fig. 7I). Approximately 75-80% of DEGs were associated with differential chromatin loops in both WT and ko2 cells, indicating a strong coupling between transcriptional changes and chromatin loop remodeling. Consistent with this observation, DEGs were significantly enriched among genes associated with both WT-specific loops (odds ratio = 1.64, Fisher’s exact test, P = 3.6 × 10 ³) and ko2-specific loops (odds ratio = 1.72, P = 2.6 × 10 ¹) (Fig. 7I). Together, these findings indicate that genes undergoing transcriptional changes are preferentially associated with regions of chromatin loop rewiring, supporting a close link between BORIS-dependent transcriptional reprogramming and three-dimensional genome organization.

## Discussion

High-grade serous ovarian carcinoma, the most common subtype of ovarian cancer, remains one of the deadliest gynecological malignancies because of its late-stage diagnosis, marked molecular heterogeneity, frequent therapeutic resistance, extensive genomic instability, and widespread epigenetic dysregulation (59). The lack of reliable biomarkers for early detection highlights the need to identify new molecular drivers and pathways involved in disease initiation and progression. In this study, we show that the germline-specific gene CTCFL (BORIS) is frequently aberrantly activated in ovarian cancer, where it reshapes chromatin architecture and establishes a distinct transcriptional program.

Cell type–specific transcriptional programs are maintained by the three-dimensional organization of the genome, which is determined by chromatin accessibility, epigenetic modifications, and CTCF-mediated chromatin architecture (4). CTCF binding sites serve as architectural anchors by blocking cohesin-mediated loop extrusion through interactions involving its N-terminal domain (12, 13). BORIS recognizes the same genomic binding sites as CTCF through its almost identical 11-zinc-finger DNA-binding domain but, unlike CTCF, lacks the ability to block cohesin-mediated loop extrusion efficiently (12). We propose that this fundamental functional difference underlies the architectural role of BORIS in cancer.

CTCF and BORIS are physiologically co-expressed during spermatogenesis, with BORIS highest expression occurring in type B spermatogonia (26), where chromatin adopts a more permissive three-dimensional organization that supports extensive transcriptional reprogramming preceding meiosis (60). A similarly permissive chromatin state has been described in many cancers, facilitating activation of oncogenic transcriptional programs (61). Our findings suggest that BORIS may contribute to the establishment of this permissive chromatin state. Consistent with this concept, BORIS depletion in OVCAR8 cells produced the opposite phenotype: increased A/B compartment segregation, stronger TAD insulation, enhanced chromatin loop strength, and widespread transcriptional reprogramming. Together, these findings support a model in which aberrant BORIS activation weakens CTCF-mediated chromatin insulation, thereby generating a more permissive chromatin architecture that supports the ovarian cancer transcriptional program.

Our study provides the first evidence that BORIS binding at CTCF-bound chromatin anchors influences genome organization at multiple hierarchical levels. Previous work demonstrated that BORIS promotes de novo chromatin interactions and super-enhancer formation in treatment-resistant neuroblastoma, contributing to therapeutic resistance (27). Our findings substantially extend these observations by showing that BORIS regulates not only chromatin looping but also compartment segregation, TAD insulation, and genome-wide transcriptional programs. Together, these studies identify BORIS as a broader regulator of CTCF-mediated three-dimensional genome architecture in cancer.

BORIS is frequently activated in a wide range of human cancers, where it has been associated with increased invasiveness (45), epithelial–mesenchymal transition (62), resistance to anticancer therapies (27), activation of cancer-testis genes (63), and regulation of human-specific transposable elements (28). Consistent with an oncogenic role, ectopic BORIS expression in somatic tissues of humanized BORIS transgenic mice increases tumor incidence (42), suggesting that aberrant BORIS activation is sufficient to promote tumorigenesis. Likewise, depletion of BORIS in several BORIS-positive cancer cell lines has been reported to induce cell death, differentiation, or a less tumorigenic phenotype (25). Our findings add another layer to the functional role of BORIS in cancer. In contrast, BORIS knockout in OVCAR8 cells did not diminish tumorigenic characteristics. Instead, BORIS-deficient cells displayed increased proliferation, enhanced drug resistance, greater migratory capacity, and a transition from a compact epithelial morphology to a more elongated mesenchymal-like phenotype. Interestingly, these findings are consistent with a previous study reporting growth-inhibitory effects of BORIS both in vitro and in vivo (64). Collectively, these observations indicate that the biological consequences of BORIS activation are highly context dependent rather than uniformly oncogenic or tumor suppressive. One possible explanation for this dependence on context is the highly rearranged genome of OVCAR8 cells. In a genome containing extensive structural abnormalities, widespread epigenetic and transcriptional remodeling following BORIS depletion may not restore a normal chromatin state but instead reinforce alternative regulatory programs that favor a more aggressive phenotype. We also cannot exclude the possibility that low-abundance BORIS isoforms lacking the deleted exons are expressed and retain partial functions independent of chromatin binding, although this remains to be experimentally addressed. More generally, the outcome of BORIS activation is likely determined by the cellular environment, including chromatin organization, DNA methylation, and the availability of interacting protein partners.

One of the major findings of this study is that BORIS loss destabilizes CTCF/cohesin occupancy and remodels the chromatin landscape. Conversely, ectopic BORIS expression in BORIS-negative somatic cells stabilizes CTCF occupancy and induces epigenetic remodeling at CTCF-bound regions (24). Together, these reciprocal observations indicate that BORIS directly modulates the chromatin environment surrounding CTCF-bound sites.

Importantly, chromatin remodeling following BORIS depletion was not entirely deterministic. Although both ko clones carried the same homozygous deletion of BORIS gene and displayed highly similar global transcriptional and chromatin phenotypes, a subset of altered CTCF/cohesin binding sites differed between the clones. These clone-specific changes likely reflect the intrinsic plasticity of the epigenome and the influence of local chromatin context during clonal expansion. In contrast, a substantial subset of altered CTCF/cohesin binding sites and transcriptional changes was shared between both clones and localized to BORIS-bound regions, representing reproducible, gene-specific consequences of BORIS loss. Thus, BORIS depletion induces both deterministic and stochastic remodeling of genome organization, with the shared alterations defining the core architectural and transcriptional functions of BORIS in ovarian cancer cells.

In summary, our findings identify BORIS as a previously unrecognized regulator of three-dimensional genome architecture in ovarian cancer. Rather than functioning as a conventional architectural protein, BORIS appears to act as a permissive chromatin anchor that occupies CTCF binding sites without efficiently blocking cohesin-mediated loop extrusion. This property weakens chromatin insulation, promotes a more permissive three-dimensional genome organization, and facilitates widespread transcriptional reprogramming. These findings establish BORIS as a clinically relevant regulator of ovarian cancer biology and support its further investigation as both a prognostic biomarker and a potential therapeutic target.

## Data availability

All raw and processed sequencing data used in this study are available through NCBI GEO database (GSE341599: https://www.ncbi.nlm.nih.gov/gds/?term=GSE341599).

## Supplementary data

Supplementary Fig. S1-S10.

## Acknowledgments

This research was supported by the Intramural Research Program of the National Institutes of Health (NIH). The contributions of the NIH authors are considered Works of the United States Government. The findings and conclusions presented in this paper are those of the authors and do not necessarily reflect the views of the NIH or the U.S. Department of Health and Human Services. This study used the Office of Cyber Infrastructure and Computational Biology High-Performance Computing cluster at NIAID, and high-performance computational capabilities of the Biowulf Linux cluster at NIH. We thank the Integrated Data Sciences Section (IDSS), especially Dr. Justin Lack, for facilitating the collaboration that provided bioinformatics support for this study. The authors acknowledge the use of ChatGPT (OpenAI) to assist with language editing and improving the clarity and flow of this manuscript. All AI-generated suggestions were verified and edited by the authors, who take full responsibility for the accuracy and final content of the publication.

## Author Contributions

D.N.B., E.M.P., L.K., D.L., M.T., and J.B. conducted the experiments. Y.J. and E.P. coordinated the ordering of reagents. D.N.B, A.S., T.M., J.L., and E.M.P. performed the bioinformatics analyses. D.N.B., V.V.L., and E.M.P. conceived and designed the project. E.M.P. wrote the paper with contributions and corrections from D.N.B., A.S., L.K., M.T. All authors reviewed and approved the manuscript.

## Funding

This work was supported by the Division of Intramural Research/NIAID/NIH (to V.V.L.).

## Conflict of interest statement

No potential conflict of interest was reported by the authors.

## Notes

### Competing Interest Statement

The authors have declared no competing interest.

