## Supporting Data for "The CTCF Paralog BORIS Contributes to the Ovarian Cancer Transcriptional Program by Relaxing CTCF-mediated 3D Genome Organization"

### Supplementary Data

#### SI Materials and Methods

##### Generation of OVCAR8 BORIS knockout cell lines.

OVCAR8 cell line was obtained from NCI collection of NCI-60 cancer cell lines. The cell line was maintained in RPMI 1640 supplemented with 10% FBS and 1% penicillin/streptomycin. BORIS/CTCFL knockout OVCAR8 cell lines were generated using CRISPR/Cas9 ribonucleoprotein (RNP)-mediated genome editing. OVCAR8 cells in logarithmic growth phase were harvested, washed with PBS, and $1 \times 10^6$ cells were collected for electroporation. Two guide RNAs (gRNAs) targeting the BORIS locus were used: gRNA-1, 5′-ATCTGCCCTGTTCAGTACGGTGG-3′, and gRNA-2, 5′-GTACCGCCAAACCTGTTAGAAGG-3′. Each gRNA (10 pmol) was separately complexed with 10 pmol of recombinant Streptococcus pyogenes Cas9 nuclease (HyCyte™ Starfish Cas9 Nuclease Multi-NLS; Product Code GUCN-R003) to generate Cas9 RNP complexes. Electroporation was performed using the Neon™ Transfection System (Thermo Fisher Scientific) according to the manufacturer's instructions. Briefly, cell pellets were resuspended in 10 μL of R Buffer containing the Cas9 RNP complexes and electroporated using the following parameters: 1,500 V, 30 ms pulse width, and one pulse. Immediately after electroporation, cells were transferred into pre-warmed complete culture medium and allowed to recover under standard culture conditions. Following recovery, cells were single-cell cloned by limiting dilution to establish monoclonal cell lines. Genomic DNA from individual clones was screened by PCR using primers flanking the targeted region (forward: 5′-AACCAGATCCCCCAAGGGTTA-3′; reverse: 5′-GAAACCCGGCTGCACTGTTA-3′), and the resulting PCR products were analyzed by Sanger sequencing to confirm CRISPR/Cas9-mediated deletion of the targeted BORIS genomic region. Clones harboring the expected deletion were selected as BORIS knockout cell lines.

##### Cell proliferation assay:

Cell proliferation was assessed using an MTT colorimetric assay kit (Cat. No. 10009365; Cayman Chemical) according to the manufacturer's protocol. Briefly, WT and ko OVCAR8 cells were seeded in sterile 96-well plates at a density of $1 \times 10^3$ cells per well in 100 μL complete culture medium. Cells were maintained at 37°C in a humidified incubator containing 5% $CO_2$ for five consecutive days. After each day incubation, 10 μL of MTT working solution was added to each well and mixed gently. Plates were then incubated for 4 hours at 37°C to allow formazan crystal formation. Next, 100 μL of freshly prepared crystal-dissolving solution was added to each well, and the plates were incubated at 37°C for 18 hours until the formazan crystals were completely dissolved. Absorbance was measured at 570 nm using a Cytation C10 confocal imaging reader (BioTek). Cell proliferation was determined based on the absorbance values relative to the control group.

##### Cell cycle analysis

Cell cycle distribution was analyzed using the BD Pharmingen™ 647 EdU Click Proliferation Kit (Cat. No. 565456, BD Biosciences) according to the manufacturer's instructions. Wild-type and BORIS knockout OVCAR8 cells were seeded in 100-mm culture dishes and incubated overnight to achieve 60-70% confluence. Cells were labeled with 10 μM 5-ethynyl-2′-

deoxyuridine (EdU) for 1 hour at 37°C in a humidified incubator with 5% $CO_2$. Following EdU incorporation, cells were washed with phosphate-buffered saline (PBS), detached using trypsin-EDTA, collected by centrifugation, and resuspended in staining buffer. Approximately $1 \times 10^6$ cells per sample were fixed with 4% paraformaldehyde-based fixative solution for 15 min at room temperature, permeabilized with 1× saponin-based permeabilization buffer, and subjected to the click chemistry reaction using Eterneon™ Red 645 azide for 30 min in the dark. After two washes, cells were stained with DAPI for total DNA content. Flow cytometric analysis was performed with BD FACSymphony A5 to determine cell cycle distribution based on EdU incorporation and DNA content. Unstained cells, DAPI-only controls, and EdU-negative controls were included to establish gating and compensation. Three independent biological replicates were analyzed for each experimental group. Finally, obtain data were analyzed using FCS Express software.

**Microscopy**

Immunofluorescent cells were imaged with a Zeiss LSM 880 confocal microscope using a plan apochromat 20X, 0.8 NA objective lens. Cells were excited in two separate channels with 405 nm and 488 nm lasers, and fluorescence emission was detected using 410 - 495 nm and 495 - 574 nm bandpass windows respectively for DAPI and Alexa 488 dyes.

**Colony formation assay**

Wild-type and BORIS knockout OVCAR8 cells were seeded at densities of 1,000 or 2,000 cells per 10-cm dish and cultured for 14 days to allow colony formation. Colonies were fixed with cold acetic acid: methanol fixing solution (1:7) for 10 min on ice, stained with 0.5% crystal violet, imaged, and quantified using ImageJ. Colony number and staining intensity were measured for each condition.

**Chemosensitivity assay**

Drug response to carboplatin and cisplatin was evaluated using an MTT-based cell viability assay. WT and ko cells were plated in 96-well plates at a density of $1 \times 10^4$ cells per well and allowed to adhere overnight at 37°C in a humidified incubator with 5% $CO_2$. Cells were then exposed to increasing concentrations of carboplatin (Millipore-Sigma, catalog #BP711) or cisplatin (USP, catalog #1134357) for 72 hours. Following treatment, MTT reagent was added to each well and incubated for 4 hours at 37°C. The resulting formazan crystals were solubilized using the crystal-dissolving solution, and absorbance was recorded at 570 nm using a Cytation C10 confocal imaging reader (BioTek).

**Scratch wound-healing assay**

Wild-type (WT) and BORIS knockout (ko) OVCAR8 cells were seeded into 24-well plates and grown to a confluent monolayer. Uniform linear wounds were generated using the AutoScratch wound-making tool (BioTek). Detached cells and debris were removed by gently washing the wells with phosphate-buffered saline (PBS), after which fresh complete culture medium was added. The plates were incubated in a Cytation 5 cell imaging multi-mode reader (BioTek) at 37°C with 5% $CO_2$ for 24 hours, and images of the cell-free area were acquired at multiple time points. Wound closure was quantified using ImageJ by measuring the remaining wound area at each time point and calculating the percentage of wound closure relative to the initial wound area.

**Western blotting**

Whole-cell protein extracts were prepared in RIPA lysis buffer (Millipore; 50 mM Tris-HCl, pH 7.4, 1% Nonidet P-40, 0.25% sodium deoxycholate, 500 mM NaCl, 1 mM EDTA) supplemented with protease inhibitor cocktail (Roche). Proteins were separated by SDS-PAGE, transferred to PVDF membranes, and immunoblotted with rabbit recombinant monoclonal anti-BORIS antibody (Abcam, EP12204). Rabbit monoclonal anti-histone H3 antibody (Abcam, ab201456) was used as a loading control.

**ChIP-seq**

For ChIP-seq, $2 \times 10^6$ asynchronously growing cells were crosslinked with 1% formaldehyde for 10 min at room temperature, followed by quenching with 125 mM glycine for 10 min. Cells were washed twice with phosphate-buffered saline (PBS) and lysed in ChIP lysis buffer (150 mM NaCl, 1% Triton X-100, 0.1% SDS, 20 mM Tris-HCl, pH 8.0, and 2 mM EDTA). Chromatin was sheared to an average fragment size of 200–600 bp and immunoprecipitated using 5 µg of the indicated antibody (rabbit recombinant monoclonal BORIS antibody (Abcam, EP12204); rabbit polyclonal antibody against histone H2A.Z (Abcam, ab4174); mouse monoclonal antibody against CTCF (Santa Cruz, (B-5): sc-271514); rabbit polyclonal to Histone H3 (acetyl K27) (Abcam, ab4729); rabbit polyclonal to Histone H3 (tri methyl K4) (Abcam, ab8580), rabbit monoclonal antibody against RAD21 (Abcam, ab217678), rabbit polyclonal antibody against SMC3 (Abcam, ab#9263). Immunoprecipitated DNA was purified using ChIP DNA Clean & Concentrator (Zima Research, D5205). DNA concentration was quantified using a Qubit 4 Fluorometer (Thermo Fisher Scientific), and 5–10 ng of DNA was used for library preparation with the NEBNext Ultra II DNA Library Prep Kit for Illumina (New England Biolabs, E7645S). Libraries were sequenced using paired-end reads on an Illumina NovaSeq 6000 platform.

**RNA-seq**

Total RNA was extracted from OVCAR8 cells using TRIzol reagent according to the manufacturer's protocol. RNA-seq libraries were prepared according to the manufacturer's protocol using the NEBNext Ultra II Directional RNA Library Prep Kit for Illumina (New England Biolabs, E7760S). RNA-Seq libraries from at least three biological and technical replicates for each cell type and condition were paired-end sequenced on an Illumina NovaSeq 6000 platform.

**ATAC-seq**

ATAC-seq libraries were prepared from 50,000 OVCAR8 cells. Nuclei were isolated by lysis in cold buffer (10 mM Tris-HCl, pH 7.4, 10 mM NaCl, 3 mM MgCl_2, and 0.1% IGEPAL CA-630) and subjected to tagmentation using Tn5 transposase (Illumina) for 40 min at 37°C. Tagmented DNA was purified using DNA Clean & Concentrator-5 columns (Zymo Research), amplified with NEBNext High-Fidelity 2× PCR Master Mix (New England Biolabs), and size-selected using 1.8× AMPure XP beads (Beckman Coulter). Library concentration and quality were assessed using a Qubit fluorometer (Thermo Fisher Scientific) and an Agilent Bioanalyzer, respectively. Libraries from three biological replicates per condition were paired-end sequenced on an Illumina NovaSeq 6000 platform.

**Bioinformatic analysis of NGS data**

Sequencing reads from ChIP-seq, RNA-seq, and ATAC-seq experiments were aligned to the human reference genome (hg38). ChIP-seq and ATAC-seq reads were aligned using Bowtie2 (1), whereas RNA-seq reads were aligned using STAR with the default parameters (2). Peak calling for ChIP-seq and ATAC-seq was performed with MACS3 (3) after removal of duplicate reads, and normalized signal tracks (bigWig) were generated using DeepTools (4) for visualization in the Integrative Genomics Viewer (IGV) (5). Peak overlaps and genomic feature analyses were performed using BEDTools (6). Heatmaps and average signal profiles were generated with DeepTools (4), and de novo motif analysis was performed using the MEME Suite (7). Differential gene expression analysis was performed using DESeq2 (8). Genes with an adjusted P value (FDR) $< 0.0001$ and $|\log2 \text{ fold change}| > 1$ were considered significantly differentially expressed. Gene set enrichment analysis was performed using the DAVID (The Database for Annotation, Visualization, and Integrated Discovery) (9). Reproducibility among biological replicates was assessed by Pearson correlation analysis.

**scRNA-seq data analysis**

Publicly available datasets for both female fetal gonadal cells and high grade serious ovarian cancer samples were downloaded from Gene Expression Omnibus (GEO). The fetal gonad dataset (GSE181558) was comprised female gonad mesonephros and fallopian tube tissue from 11 donors aged 6-18 WPF (10). The HGSOC (GSE154600) samples were obtained from tumor specimens that were used for clinical diagnoses of ovarian cancer; the excess was used in these studies (11). Both the fetal dataset and HGSOC dataset were processed using the 10x Genomics platform. The datasets were processed according to a standard Scanpy pipeline (12). Raw counts matrices were imported as AnnData objects and underwent quality control filtering for low quality cells and low expressed genes. The cells were clustered using the Leiden algorithm, and clusters were annotated using know markers (10, 11) and the database CellMarker 2.0 (13). Cell types were predicted using Python's decoupler module, using the univariate linear model (ULM) enrichment analysis pipeline. Each cell was initially assigned a cell type based on ULM enrichment scores, and then cluster-level assignments were made by ranking enrichment scores across the Leiden cluster. Final predictions were manually curated by comparing differential gene expression across clusters using Scanpy's rank_gene_groups function to identify marker genes, and results were visualized using UMAP. For the single cell binary UMAPs: the log1p normalized expression values were used. For each gene, cells with expression values greater than 0.1 were classified as positive, and cells with expression values less than or equal to 0.1 were classified as negative.

**The sources of downloaded data**

The expression of BORIS in human cancer was downloaded from GEPIA (14). The survival plot of 483 patients with ovarian cancer was downloaded from GSE14764 (15). For the analysis of gene expression in ovarian cancer (Figure 1C), we downloaded RNA-seq data from TCGA (The Cancer Genome Atlas Pan-Cancer analysis project) (16): eleven BORIS-highly positive ovarian cancers (TCGA24146901, TCGA59A5PD01, TCGA29242801, TCGA24229001, TCGA36157501, TCGA24202701, TCGA20168301, TCGA23112201, TCGA24229701, TCGA24228101, TCGA04136201) and eleven BORIS negative ovarian cancers (TCGA09036701, TCGA20168701, TCGA23102301, TCGA24147401, TCGA24154401, TCGA24184701, TCGA24202001, TCGA25132801, TCGA30189101, TCGA36157801, TCGA61172101). BORIS expression at different stages of ovarian cancer was downloaded from GENT2 platform (17). The CNV frequencies for different cancers were downloaded from cBioPortal (Cancer Genomics Data Server) (18). The expression of BORIS in normal fallopian tube epithelium, precursor lesions, and high-grade serous carcinoma using NanoString GeoMx DSP data were downloaded from (19). The Kaplan–Meier analysis of ovarian cancer were downloaded from https://kmplot.com/ (20).

**Identification of Differentially Expressed Gene Clusters.**

To identify genomic clusters of coordinately regulated genes, differentially expressed genes were analyzed based on their genomic positions. Genes were ordered by chromosome and genomic coordinate, and clusters were defined as two or more consecutive DEGs on the same chromosome separated by less than 100 kb. Upregulated and downregulated DEGs were analyzed independently. Statistical significance of clustering was assessed by permutation testing as previously described (21), in which observed cluster metrics were compared with an empirical null distribution generated by random permutation of gene positions. Clusters with $P < 0.005$ were considered statistically significant.

**Hi-C library preparation and data processing**

Hi-C libraries were prepared using the Arima-HiC Kit (Arima Genomics) according to the manufacturer's instructions. Briefly, $5 \times 10^6$ OVCAR8 cells were crosslinked with 2% formaldehyde in growth medium for 10 min at room temperature, and the reaction was quenched with 0.125 M glycine for 10 min. Cells were washed twice with phosphate-buffered saline (PBS), snap-frozen, and stored at −80°C until nuclei isolation. Hi-C data from wild-type (WT) and BORIS knockout (KO) OVCAR8 cells (two biological replicates per condition) were processed using the Juicer pipeline (22) and aligned to the human reference genome (hg38) using BWA-MEM (23). Hi-C pair files (.pairs.gz) were generated from the aligned BAM files using Pairtools (24) with default parameters. The .pairs.gz files were converted into .hic contact matrix files using Juicer. Contact matrices were normalized using the Knight–Ruiz (KR) algorithm and converted to the multi-resolution .mcool format with Cooler package (25) for downstream analyses. Downstream Hi-C analyses and visualization, including compartment analysis, saddle plots, aggregate contact analyses, and TAD visualization, were performed using the GENOVA package (26).

**TAD boundary analysis**

Topologically associating domains (TADs) were initially identified using Arrowhead (Juicer Tools v3.0.0) (22). For quantitative comparison of boundary strength between conditions, insulation scores were calculated using Cooltools (v0.7.1) (27) at 10-kb resolution with a 200-kb sliding window (--ignore-diags 2). Boundary bins were defined based on the insulation boundary flag and the corresponding boundary strength metric. To enable replicate-aware comparisons, boundary bins from all samples were merged within ±20 kb to generate a union boundary set. For each replicate, the mean boundary strength was quantified across each union boundary region using BEDTools. Differential boundary strength was calculated as the mean boundary strength in BORIS knockout cells minus the mean boundary strength in wild-type cells. Boundaries were classified as lost, gained, or shared according to their presence in both replicates of each condition. Global differences in boundary strength were evaluated using a paired Wilcoxon signed-rank test across all union boundaries, whereas statistical significance for individual boundaries was assessed using Welch's two-sample t-test. Because only two biological replicates were available per condition, the magnitude of the change in boundary strength (Δ boundary strength) was emphasized as the primary measure of biological significance. Volcano plots and scatter plots were generated in Python using Matplotlib.

**Chromatin loop identification and integration**

Chromatin loops were identified from Hi-C .hic files using Mustache (28). Loop calling was performed independently for each biological replicate at multiple resolutions (5, 10, 25, and 50 kb) to capture chromatin interactions across different spatial scales. The following parameters were applied uniformly across all samples: -pt 0.05 (candidate loop P-value threshold), -st 0.7 (sparsity threshold), and -sz 1 (minimum detection scale). Candidate loops were assigned P values using Mustache's local background model and corrected for multiple testing using the Benjamini–Hochberg procedure. Loops with a false discovery rate (FDR) < 0.05 were retained for downstream analyses. To generate non-redundant loop sets, loop calls identified at different resolutions were merged by clustering loops with anchor coordinates located within a distance tolerance corresponding to the calling resolution. Within each cluster, a representative loop was selected based on the highest spatial resolution (smallest bin size) and, when applicable, the lowest FDR. Consensus loop anchors were defined as the median coordinates of the clustered calls. Biological replicates were subsequently merged using the same coordinate-based clustering strategy, and only loops detected in both biological replicates were considered reproducible. Identical loop-calling and integration parameters were applied to all conditions to enable unbiased comparisons of loop number and loop strength.

To characterize the architectural features of chromatin loop anchors, CTCF and BORIS ChIP-seq peaks were assigned to loop anchors using a hierarchical annotation strategy. First, CTCF and BORIS peaks overlapping by at least 1 bp were classified as CTCF/BORIS co-bound sites and assigned the highest priority. Remaining CTCF peaks that did not overlap BORIS peaks were classified as CTCF-only sites. Finally, remaining BORIS peaks that did not overlap CTCF peaks were classified as BORIS-only sites. CTCF and BORIS peak sets were derived from wild-type (WT) OVCAR8 cells, as BORIS is absent in BORIS knockout cells. Each loop anchor was assigned a single annotation according to the highest-priority feature present within the anchor region: (i) CTCF/BORIS co-bound, (ii) CTCF-only, (iii) BORIS-only, or (iv) none. Thus, anchors containing both co-bound and factor-specific peaks were classified as CTCF/BORIS co-bound. Loop classes were then defined by the combination of annotations at the two anchors (e.g., CTCF-only/CTCF-only, CTCF-only/CTCF/BORIS co-bound, and CTCF/BORIS co-bound/CTCF/BORIS co-bound). The distribution of anchor classes was compared among shared, WT-specific, and ko-specific loops.

**Compartmentalization analysis**

A/B compartments were identified using Cooltools. Hi-C contact matrices were converted to multi-resolution .mcool format, Knight–Ruiz (KR) balanced, and analyzed at 100-kb resolution. Genome-wide compartment scores (E1 values) were calculated using the eigs-cis function in Cooltools and subsequently oriented based on CTCF ChIP-seq profiles from wild-type OVCAR8 cells to ensure consistent assignment of A and B compartments across samples. Saddle plots were generated with GENOVA package (26) by ranking genomic bins according to their compartment scores and partitioning them into 10 equally sized quantile bins. Observed/expected contact frequencies were then aggregated for all pairwise combinations of quantile bins, generating a matrix representing interaction enrichment between compartments.

To quantify compartmentalization strength, the outer quantiles corresponding to the strongest A and B compartments (top and bottom 20% of bins) were used to calculate the mean observed/expected interaction frequencies for A–A (AA), B–B (BB), and A–B (AB) interactions. Compartment strength was calculated as the $\log_2$-transformed ratio of within-compartment to between-compartment interactions, defined as $\log_2[(AA \times BB)/(AB^2)]$, where AA and BB represent the mean interaction frequencies within A and B compartments, respectively, and AB represents the mean interaction frequency between compartments. For each sample, compartment strength was calculated for each chromosome arm and averaged to obtain a genome-wide value. For comparisons between wild-type and BORIS knockout cells, replicate values were averaged, and statistical significance was assessed using a paired Wilcoxon signed-rank test. Chromosomes 5 and 22 were excluded from the analysis because extensive structural rearrangements resulted in unreliable compartment assignments. These compartment calls were inconsistent with CTCF and H3K27ac ChIP-seq, ATAC-seq, and RNA-seq datasets and were therefore excluded from downstream analyses.

#### Supplementary Data

#### Description of Supplementary Tables:

Supplementary Table S1a: CTCF and CTCFL/BORIS expression in 300 HGSOC samples.

Supplementary Table S1b: Differentially expressed genes in BORIS-positive versus BORIS-negative HGSOC.

Supplementary Table S1c: The list of upregulated and downregulated pathways in BORIS-high HGSOC compared to BORIS low HGSOC.

Supplementary Table S1d: The list of genes with highest Sperman correlation in BORIS-high ovarian cancers.

Supplementary Table S1e: The list of genes overlapping between DEGs from S1b and genes from S1d.

Supplementary Table S2a: Differentially expressed genes in ko1 compared to WT.

Supplementary Table S2b: Differentially expressed genes in ko2 compared to WT.

Supplementary Table S2c: Differentially expressed genes in ko1/ko2 compared to WT.

Supplementary Table S2d: The list upregulated pathways in ko cells compared to WT.

Supplementary Table S2e: The list downregulated pathways in ko cells compared to WT.

Supplementary Table S3a: Differentially expressed genes deregulated in clusters in both ko1 and ko2 clones, compared to WT.

Supplementary Table S3b: Differentially expressed genes deregulated in clusters in ko1, compared to WT.

Supplementary Table S3c: Differentially expressed genes deregulated in clusters in ko2, compared to WT.

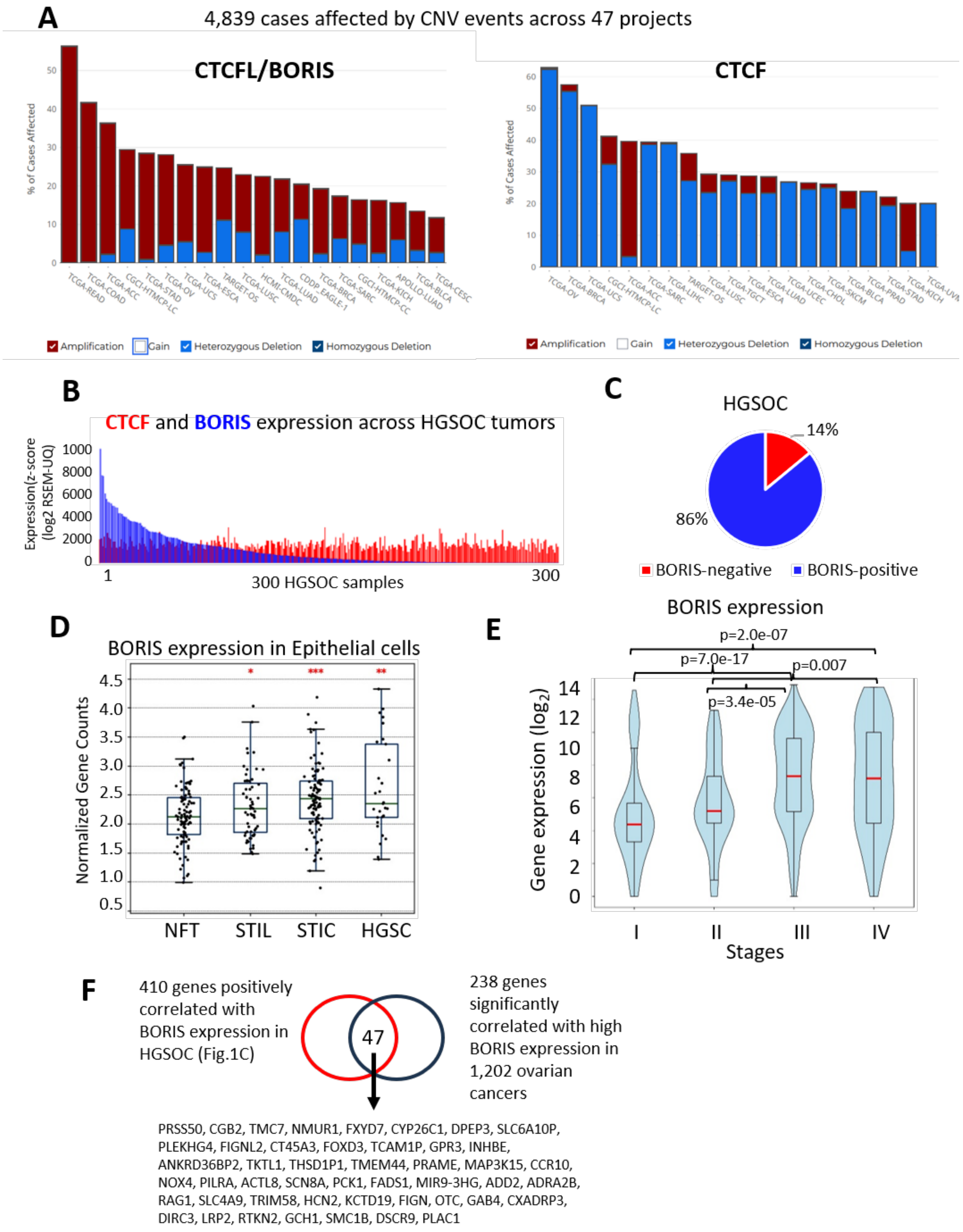
A
4,839 cases affected by CNV events across 47 projects
CTCFL/BORIS
% of Cases Affected
TCGA-READ
TCGA-COAD
TCGA-ACC
CGCI-HTMCP-LC
TCGA-STAD
TCGA-OV
TCGA-UCS
TCGA-ESCA
TARGET-OS
TCGA-LUSC
HCMI-CMDC
TCGA-LUAD
CDDP_EAGLE-1
TCGA-BRCA
TCGA-SARC
CGCI-HTMCP-CC
TCGA-KICH
APOLLO-LUAD
TCGA-BLCA
TCGA-CESC
Amplification
Gain
Heterozygous Deletion
Homozygous Deletion
CTCF
% of Cases Affected
TCGA-OV
TCGA-BRCA
TCGA-UCS
CGCI-HTMCP-LC
TCGA-ACC
TCGA-SARC
TCGA-LIHC
TARGET-OS
TCGA-LUSC
TCGA-TGCT
TCGA-ESCA
TCGA-LUAD
TCGA-UCEC
TCGA-CHOL
TCGA-SKCM
TCGA-BLCA
TCGA-PRAD
TCGA-STAD
TCGA-KICH
TCGA-UVM
Amplification
Gain
Heterozygous Deletion
Homozygous Deletion
B
CTCF and BORIS expression across HGSOC tumors
Expression(z-score (log2 RSEM-UQ)
1000
8000
6000
4000
2000
0
1
300 HGSOC samples
300
C
HGSOC
14%
86%
BORIS-negative
BORIS-positive
D
BORIS expression in Epithelial cells
Normalized Gene Counts
4.5
4.0
3.5
3.0
2.5
2.0
1.5
1.0
0.5
*
***
**
NFT
STIL
STIC
HGSC
E
BORIS expression
p=2.0e-07
p=7.0e-17
p=0.007
p=3.4e-05
Gene expression (log2)
14
12
10
8
6
4
2
0
I
II
III
IV
Stages
F
410 genes positively correlated with BORIS expression in HGSOC (Fig.1C)
47
238 genes significantly correlated with high BORIS expression in 1,202 ovarian cancers
PRSS50, CGB2, TMC7, NMUR1, FXYD7, CYP26C1, DPEP3, SLC6A10P,
PLEKHG4, FIGNL2, CT45A3, FOXD3, TCAM1P, GPR3, INHBE,
ANKRD36BP2, TKTL1, THSD1P1, TMEM44, PRAME, MAP3K15, CCR10,
NOX4, PILRA, ACTL8, SCN8A, PCK1, FADS1, MIR9-3HG, ADD2, ADRA2B,
RAG1, SLC4A9, TRIM58, HCN2, KCTD19, FIGN, OTC, GAB4, CXADRP3,
DIRC3, LRP2, RTKN2, GCH1, SMC1B, DSCR9, PLAC1

#### Supplementary Data

**Figure S1. Genomic alterations and expression of BORIS in high-grade serous ovarian carcinoma (HGSOC). (A)** Frequency of copy number alterations affecting CTCFL/BORIS (left) and CTCF (right) across 47 TCGA cancer projects. Stacked bar plots show the proportions of amplification, gain, heterozygous deletion, and homozygous deletion. **(B)** Relative expression of CTCF (red) and BORIS (blue) across 300 HGSOC tumors. Samples are ranked according to highest BORIS expression. **(C)** Proportion of HGSOC tumors expressing BORIS versus BORIS negative. Tumors with RSEM > 22 were classified as BORIS-positive. **(D)** BORIS expression in normal fallopian tube (NFT), serous tubal intraepithelial lesion (STIL), serous tubal intraepithelial carcinoma (STIC), and high-grade serous carcinoma (HGSC) epithelial cells using NanoString GeoMx DSP data. *, $p < 0.05$; **, $p< 0.01$; ***, $p< 0.001$. **(E)** Association between BORIS expression and tumor stage in ovarian cancer. **(F)** Overlap between genes significantly correlated with high BORIS expression in the HGSOC cohort analyzed in Fig. 1C and genes correlated with high BORIS expression in an independent cohort of 1,202 ovarian cancers, identifying a common set of 47 genes.

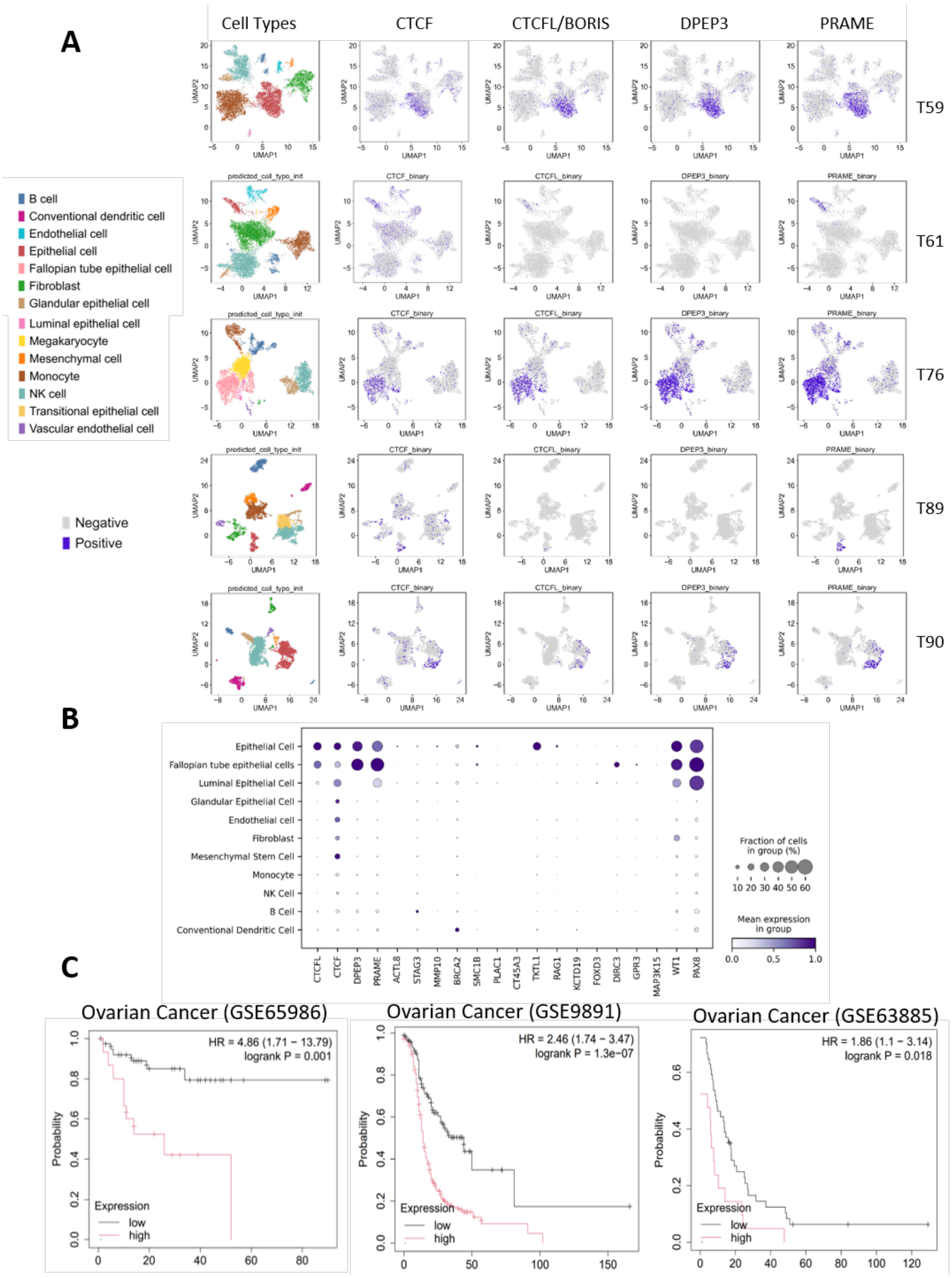
A
Cell Types
CTCF
CTCFL/BORIS
DPEP3
PRAME
T59
T61
T76
T89
T90
UMAP1
UMAP2
B cell
Conventional dendritic cell
Endothelial cell
Epithelial cell
Fallopian tube epithelial cell
Fibroblast
Glandular epithelial cell
Luminal epithelial cell
Megakaryocyte
Mesenchymal cell
Monocyte
NK cell
Transitional epithelial cell
Vascular endothelial cell
Negative
Positive
B
Epithelial Cell
Fallopian tube epithelial cells
Luminal Epithelial Cell
Glandular Epithelial Cell
Endothelial cell
Fibroblast
Mesenchymal Stem Cell
Monocyte
NK Cell
B Cell
Conventional Dendritic Cell
CTCFL
CTCF
DPEP3
PRAME
ACTL8
STAG3
MMP10
BRCA2
SMC1B
PLAC1
CT45A3
TKTL1
RAG1
KCTD19
FOXD3
DIRC3
GPR3
MAP3K15
WT1
PAX8
Fraction of cells in group (%)
10 20 30 40 50 60
Mean expression in group
0.0 0.5 1.0
C
Ovarian Cancer (GSE65986)
HR = 4.86 (1.71 − 13.79)
logrank P = 0.001
Ovarian Cancer (GSE9891)
HR = 2.46 (1.74 − 3.47)
logrank P = 1.3e−07
Ovarian Cancer (GSE63885)
HR = 1.86 (1.1 − 3.14)
logrank P = 0.018
Probability
Expression
low
high

**Figure S2. BORIS expression in HGSOC. (A)** UMAP projections of scRNA-seq datasets from five HGSOC tumors (T59, T61, T76, T89, and T90). Left panels show annotated cell types, and adjacent panels show expression of CTCF, CTCFL/BORIS, DPEP3, and PRAME. Cells expressing each gene are shown in purple**. (B)** Dot plot showing the expression of CTCFL/BORIS and selected BORIS-associated genes across annotated cell populations. Dot size represents the fraction of expressing cells, and color intensity indicates mean expression. **(C)** Kaplan–Meier survival analysis of three independent ovarian cancer cohorts (GSE65986, GSE9891, GSE63885) stratified by BORIS expression, demonstrating poorer overall survival in patients with high BORIS expression. Hazard ratios (HRs) and log-rank P values are indicated.

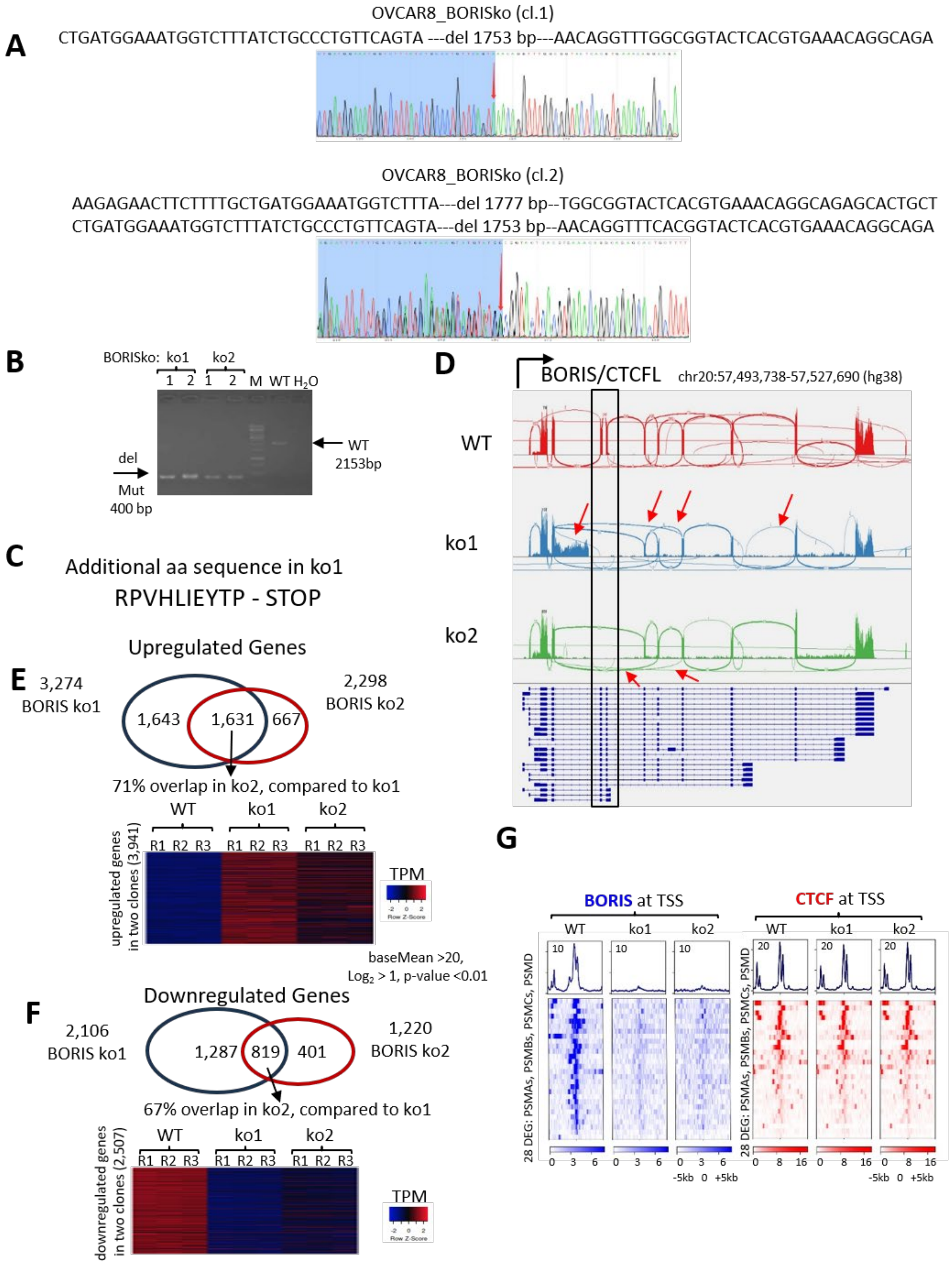
A
OVCAR8_BORISko (cl.1)
CTGATGGAAATGGTCTTTATCTGCCCTGTTCAGTA ---del 1753 bp---AACAGGTTTGGCGGTACTCACGTGAAACAGGCAGA
OVCAR8_BORISko (cl.2)
AAGAGAACTTCTTTTGCTGATGGAAATGGTCTTTA---del 1777 bp--TGGCGGTACTCACGTGAAACAGGCAGAGCACTGCT
CTGATGGAAATGGTCTTTATCTGCCCTGTTCAGTA---del 1753 bp--AACAGGTTTCACGGTACTCACGTGAAACAGGCAGA
B
BORISko: ko1 ko2
1 2 1 2 M WT H2O
del
Mut
400 bp
WT
2153bp
C
Additional aa sequence in ko1
RPVHLIEYTP - STOP
D
BORIS/CTCFL chr20:57,493,738-57,527,690 (hg38)
WT
ko1
ko2
E
Upregulated Genes
3,274
BORIS ko1
1,643
1,631
667
2,298
BORIS ko2
71% overlap in ko2, compared to ko1
WT ko1 ko2
R1 R2 R3 R1 R2 R3 R1 R2 R3
upregulated genes in two clones (3,941)
TPM
Row Z-Score
baseMean >20,
Log2 > 1, p-value <0.01
F
Downregulated Genes
2,106
BORIS ko1
1,287
819
401
1,220
BORIS ko2
67% overlap in ko2, compared to ko1
WT ko1 ko2
R1 R2 R3 R1 R2 R3 R1 R2 R3
downregulated genes in two clones (2,507)
TPM
Row Z-Score
G
BORIS at TSS
CTCF at TSS
WT ko1 ko2
28 DEG: PSMAs, PSMBs, PSMCs, PSMD
-5kb 0 +5kb

**Figure S3**. **Validation of BORIS knockout and reproducibility of transcriptional changes in OVCAR8 cells. (A)** Representative Sanger sequencing chromatograms of BORIS knockout (ko) clone 1 (top) and clone 2 (bottom). In both clones, sequencing traces show the breakpoint at the deletion junction (red arrows), confirming removal of approximately 1.7 kb of genomic DNA within the *BORIS/CTCFL* locus. **(B)** Genomic PCR analysis confirms deletion of an approximately 1.7-kb genomic fragment in both ko clones. Whereas the wild-type (WT) allele yields a 2,153-bp PCR product, both ko clones produce a ~400-bp amplicon consistent with the expected deletion. **(C)** In one allele of ko1, extension of the open reading frame (ORF) from exon 3 into the adjacent intronic sequence results in the addition of 10 novel amino acids followed by a premature stop codon. **(D)** Sashimi plot of RNA-seq data across the *BORIS/CTCFL* locus showing altered splice junction usage (red arrows) and deletion of exons 4–5 (black box) in both BORIS ko clones relative to WT. **(E,F)** Comparison of differentially expressed genes between the two independent BORIS ko clones. Venn diagrams show the overlap of significantly upregulated **(E)** and downregulated **(F)** genes. Heatmaps display genes consistently upregulated or downregulated in both koclones relative to WT. Differential expression analysis was performed using baseMean > 20, $\log_2$ fold change > 1, and $p < 0.01$. **(G)** Heatmaps and average signal profiles of BORIS and CTCF ChIP-seq signals centered on the transcription start sites (TSSs) of 28 proteasome-associated differentially expressed genes in WT and BORIS ko clones, demonstrating loss of BORIS occupancy while CTCF binding is largely maintained.

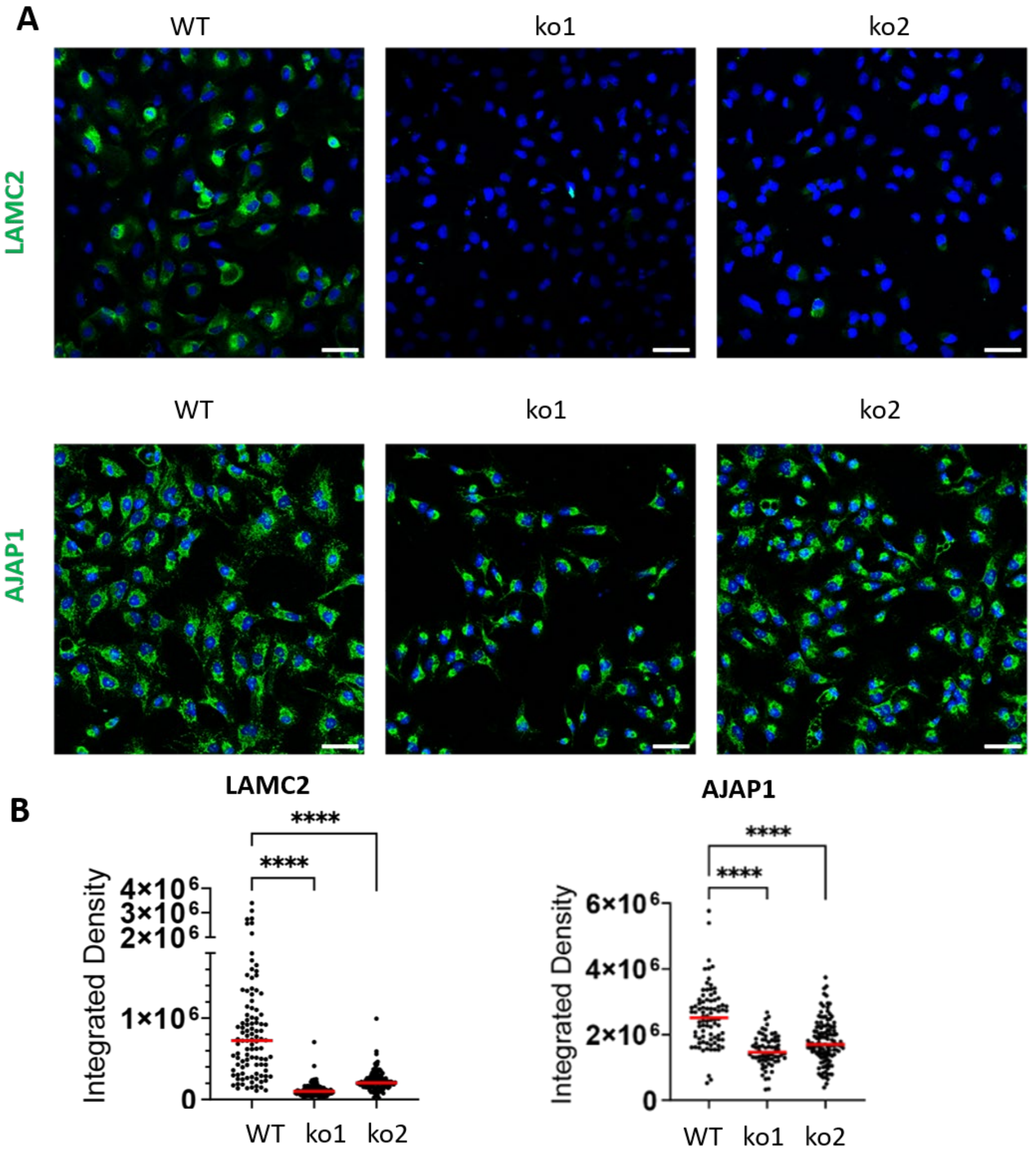


**Figure S4. Reduced AJAP1 and LAMC2 protein expression in BORIS knockout OVCAR8 cells. (A)** Representative immunofluorescence images of wild-type (WT) and two independent BORIS knockout (ko) clones stained for LAMC2 (top, green) or AJAP1 (bottom, green). Nuclei were counterstained with DAPI (blue). All AJAP1 Images and all LAMC2 images were acquired and scaled identically to allow direct signal intensity comparison. Scale bars, 50 µm**. (B)** Quantification of LAMC2 and AJAP1 fluorescence intensity. Integrated density of fluorescent signal was measured on a per-cell basis. Red lines indicate the mean. ****P < 0.0001, one-way Welch's ANOVA.

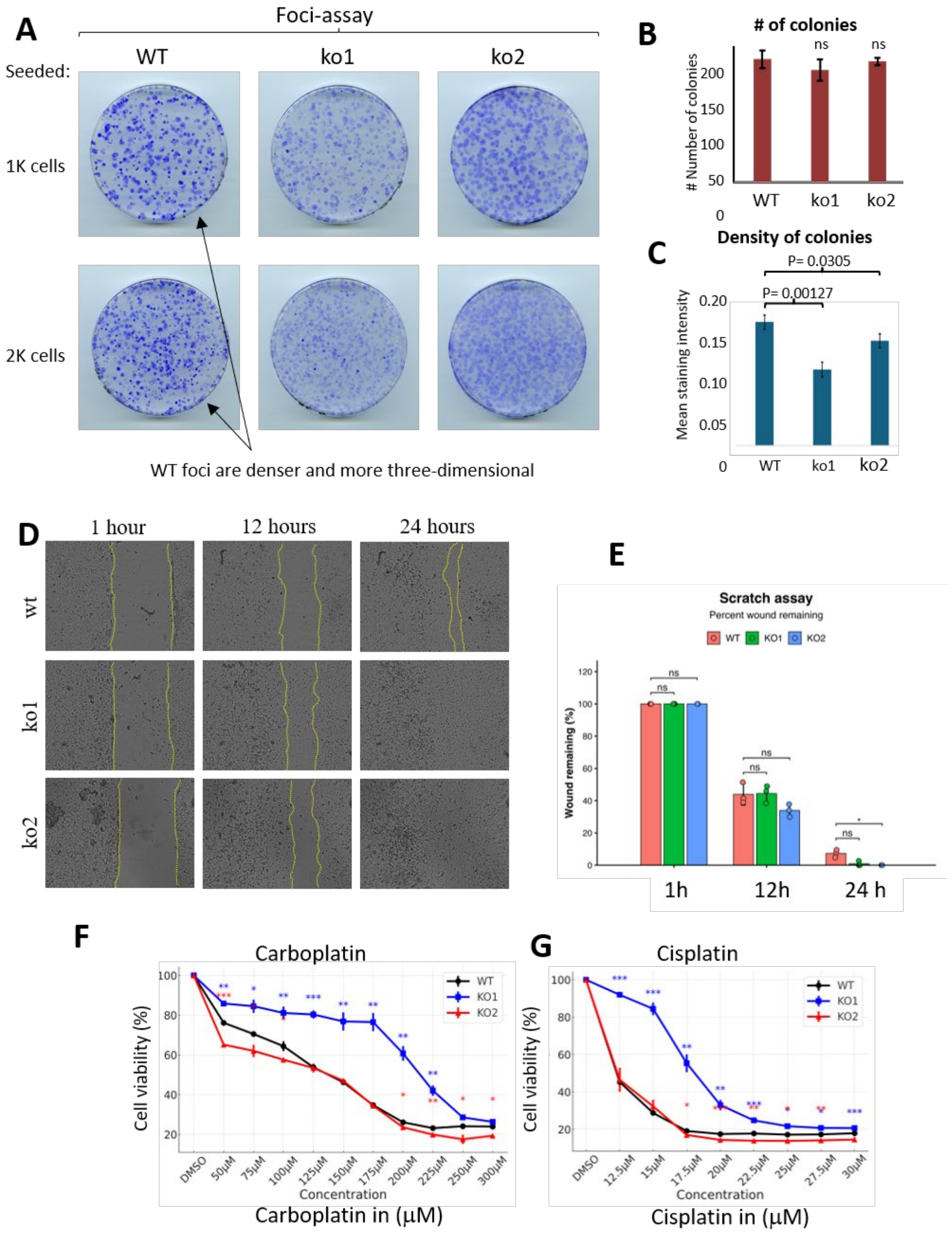


*, **, *** - *p* < 0.05, **p** < 0.01, ***p*** < 0.001 (t-test, WT vs ko1, WT vs ko2)

**Figure S5. BORIS knockout increases platinum sensitivity but does not impair clonogenic growth of OVCAR8 cells. (A)** Representative crystal violet-stained colony (foci) formation assays of WT and BORIS ko clones seeded at 1,000 or 2,000 cells per well. Although the total number of colonies was similar, WT colonies appeared denser and more compact than those formed by BORIS ko cells. **(B)** Quantification of colony number showing no significant difference between WT and BORIS ko clones. **(C)** Quantification of colony staining intensity (colony density) demonstrating reduced colony density in BORIS ko clones compared with WT. Statistical significance was determined using Welch's t-test. **(D)** Representative images from scratch (wound-healing) assays performed with WT and BORIS ko cells at 1, 12, and 24 h after scratch formation. Yellow dashed lines indicate the wound edges. **(E)** Quantification of wound closure showing significant differences in migration between WT and ko2 cells at the 24h time points. Data are presented as mean ± SEM. Paired t-test, * p-value<0.05, ns-not significant. **(F-G)** Dose-response curves showing the viability of wild-type (WT) and two independent BORIS ko clones following treatment with increasing concentrations of carboplatin **(F)** or cisplatin **(G).** Data are presented as mean ± SEM. Statistical significance was determined by unpaired two-tailed t-test ($P < 0.05$, $P < 0.01$, $P < 0.001$; WT vs. ko1, blue; WT vs. ko2, red).

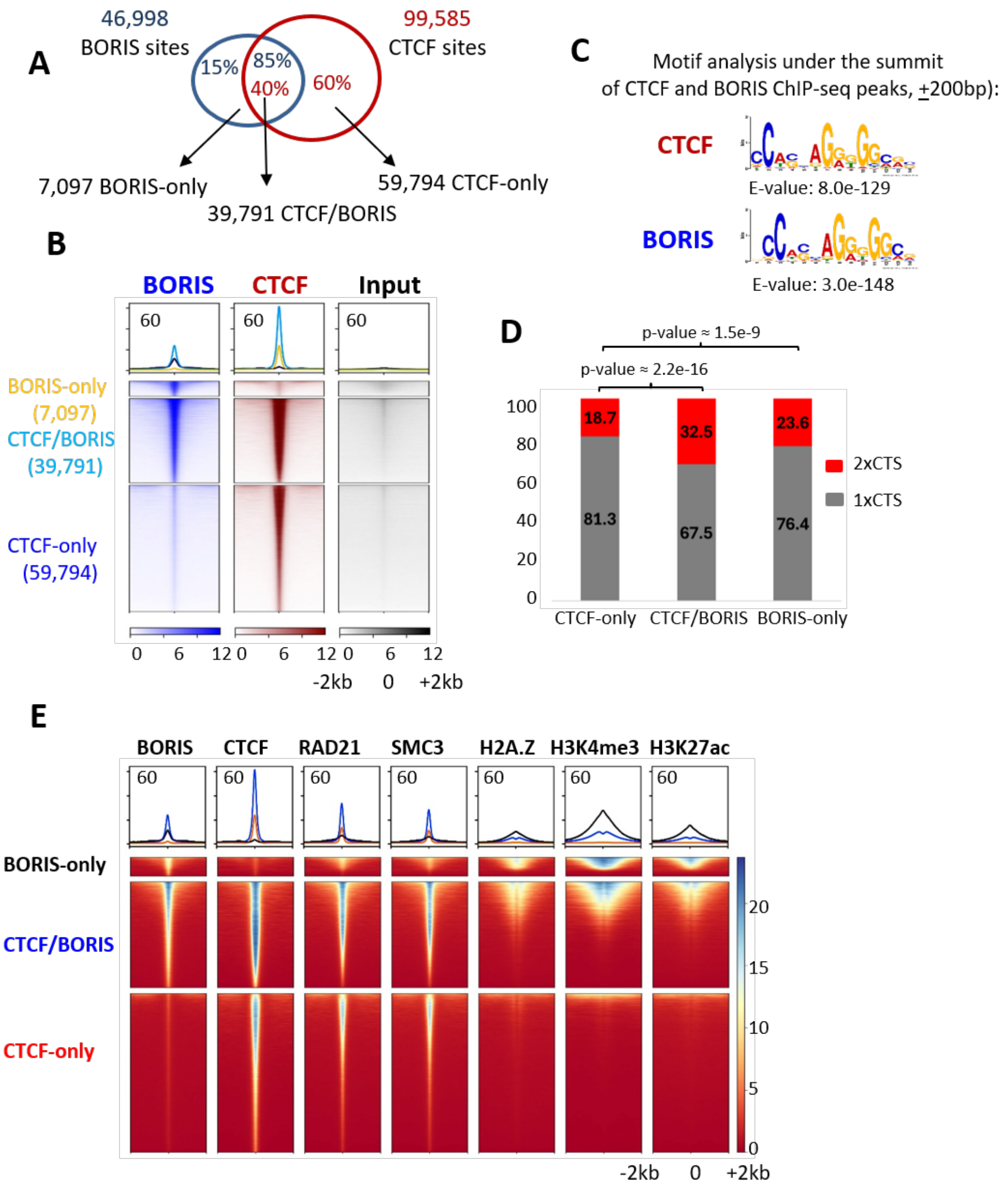


**Figure S6**. **Genomic distribution and chromatin features of BORIS and CTCF binding sites in OVCAR8 cells. (A)** Overlap between BORIS and CTCF ChIP-seq peaks. **(B)** Heatmaps and average signal profiles of BORIS, CTCF, and input ChIP-seq signals centered on BORIS-only, CTCF/BORIS, and CTCF-only binding sites (±2 kb). **(C)** De novo motif analysis of sequences underlying BORIS and CTCF ChIP-seq peak summits (±200 bp) identifies the canonical CTCF DNA-binding motif as the most significantly enriched motif in both datasets. **(D)** Proportion of genomic sites containing one (1× CTS) or two (2× CTS) CTCF-binding motifs within ±200 bp of the peak summit for CTCF-only, CTCF/BORIS, and BORIS-only peaks. Statistical significance was assessed using the chi-square test. **(E)** Heatmaps and average signal profiles of BORIS, CTCF, RAD21, SMC3, H2A.Z, H3K4me3, and H3K27ac ChIP-seq signals centered on BORIS-only, CTCF/BORIS, and CTCF-only binding sites (±2 kb).

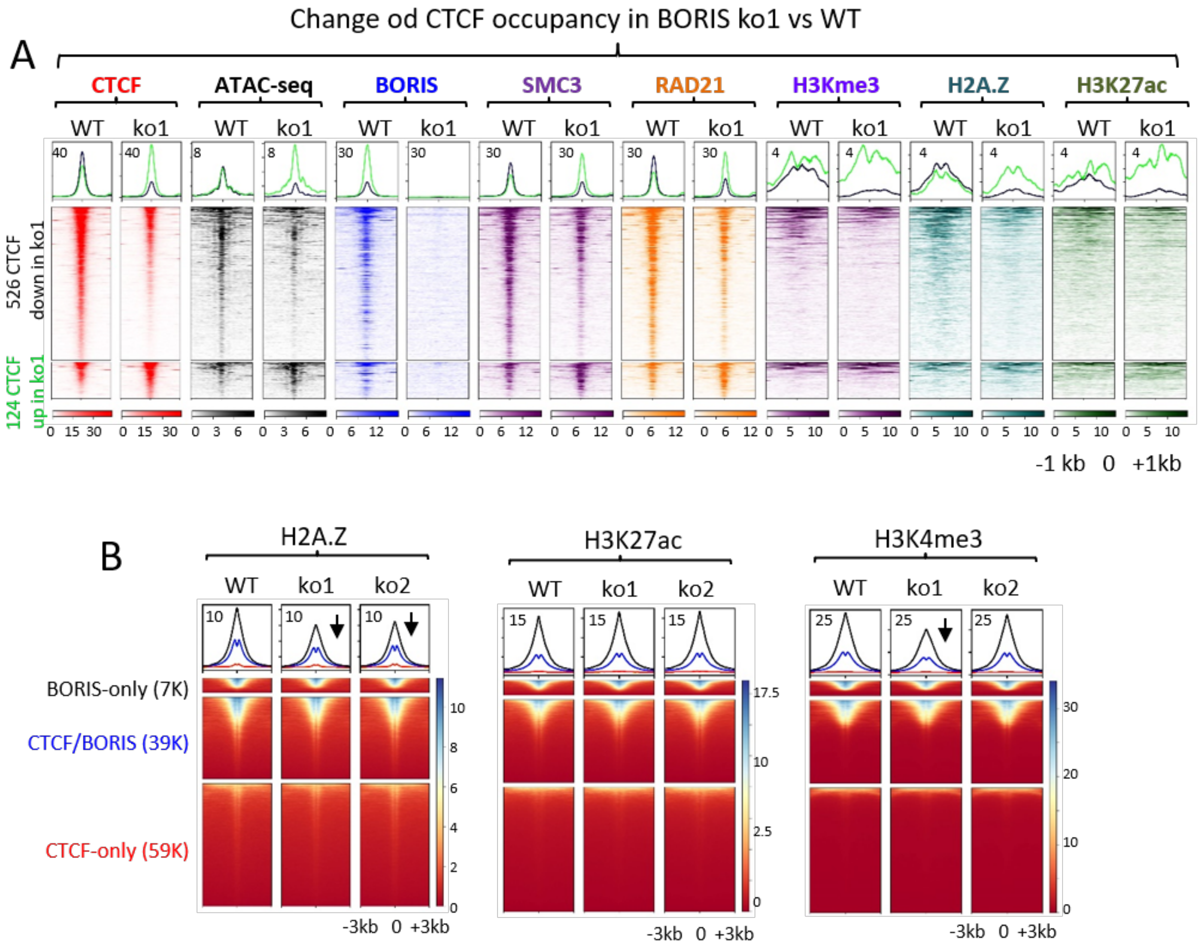


**Figure S7. Chromatin changes associated with BORIS knockout in OVCAR8. (A)** Heatmaps and average signal profiles of CTCF, ATAC-seq, BORIS, SMC3, RAD21, H3K4me3, H2A.Z, and H3K27ac signals centered on genomic regions showing increased (top, n = 526) or decreased (bottom, n = 124) CTCF occupancy in ko1 cells relative to WT. Signals are displayed within ±1 kb of the peak center. **(B)** Heatmaps and average signal profiles of H2A.Z, H3K27ac, and H3K4me3 centered on BORIS-only, CTCF/BORIS, and CTCF-only binding sites in WT and two independent BORIS knockout clones (ko1 and ko2). Signals are shown within ±3 kb of peak centers. Arrows indicate reduced enrichment of H2A.Z and H3K4me3 at BORIS-only sites following BORIS deletion.

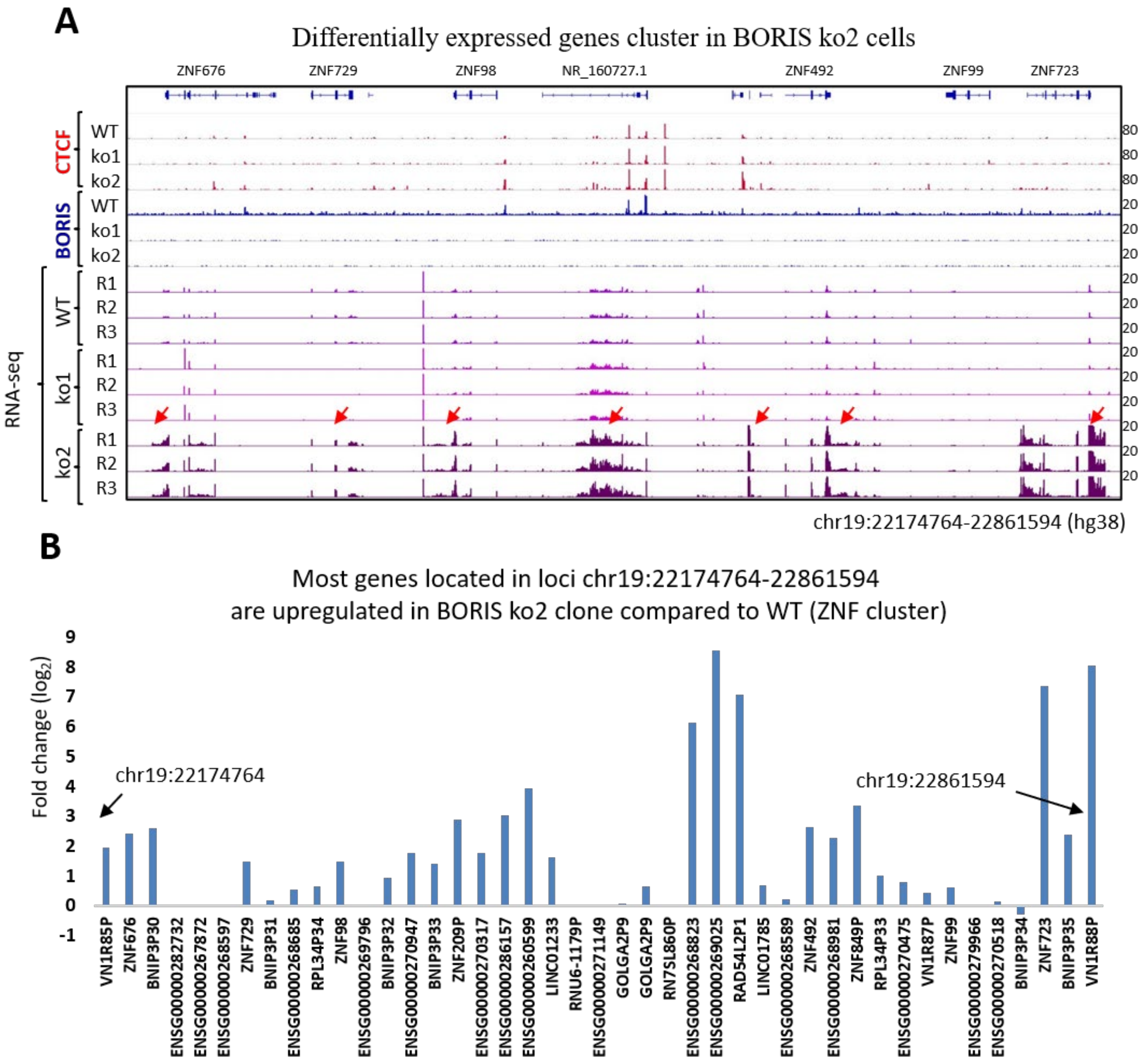


**Figure S8. Coordinated activation of a gene cluster on chromosome 19 in BORIS ko2. (A)** Genome browser view of the chromosome 19 locus showing CTCF and BORIS ChIP-seq signals and RNA-seq coverage in WT, ko1, and ko2 cells. Red arrows indicate genes displaying increased expression specifically in ko2. **(B)** $Log_2$ fold changes in expression of genes located within the same genomic interval in ko2 relative to WT, demonstrating coordinated upregulation of multiple genes across the locus.

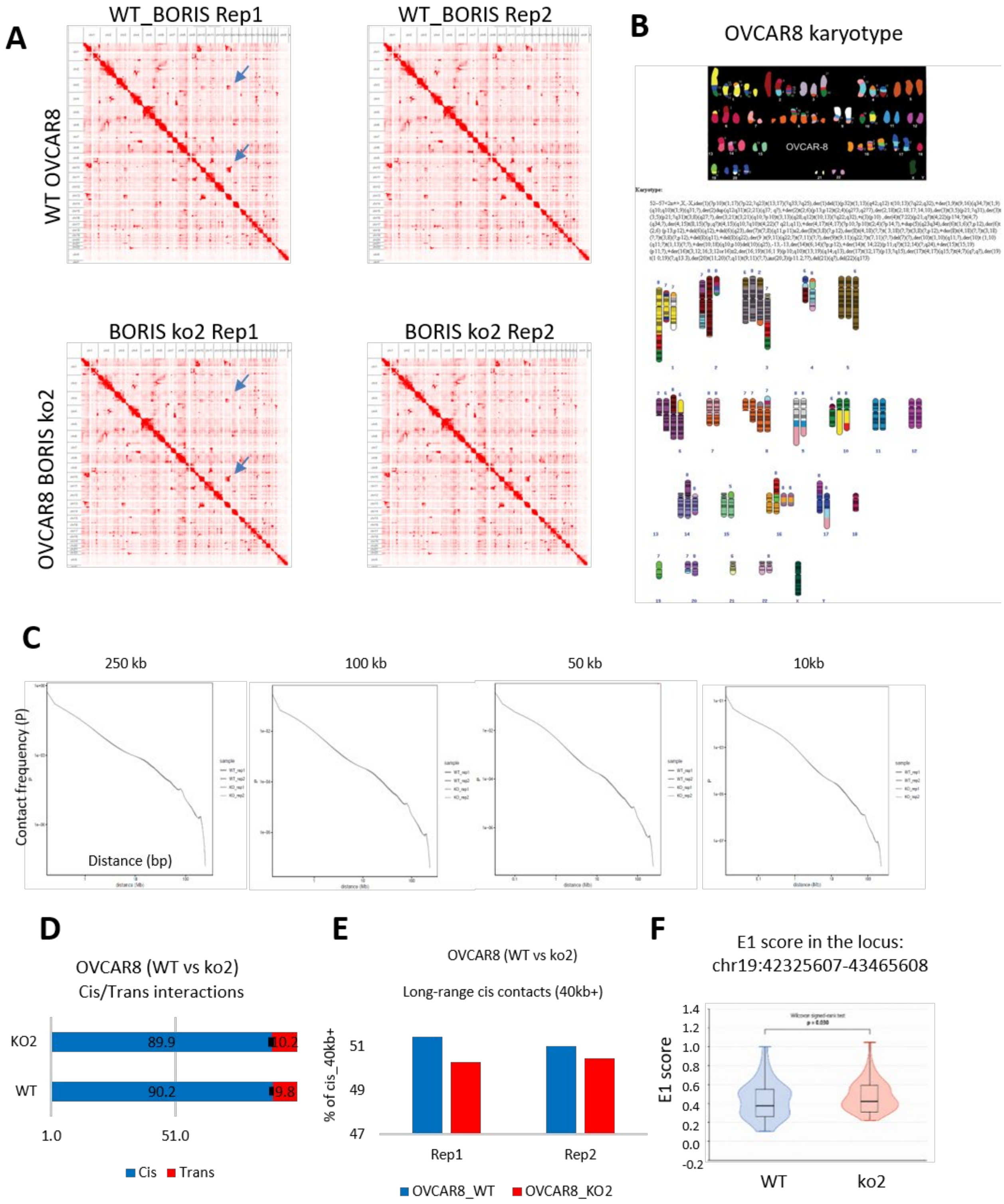
A
WT_BORIS Rep1
WT_BORIS Rep2
WT OVCAR8
BORIS ko2 Rep1
BORIS ko2 Rep2
OVCAR8 BORIS ko2
B
OVCAR8 karyotype
OVCAR-8
Karyotype:
C
250 kb
100 kb
50 kb
10kb
Contact frequency (P)
Distance (bp)
D
OVCAR8 (WT vs ko2)
Cis/Trans interactions
KO2
WT
89.9
10.2
90.2
9.8
1.0
51.0
Cis
Trans
E
OVCAR8 (WT vs ko2)
Long-range cis contacts (40kb+)
% of cis_40kb+
51
49
47
Rep1
Rep2
OVCAR8_WT
OVCAR8_KO2
F
E1 score in the locus:
chr19:42325607-43465608
E1 score
1.4
1.2
1.0
0.8
0.6
0.4
0.2
0.0
-0.2
WT
ko2

**Figure S9. Quality assessment of Hi-C data and global chromatin organization in BORIS knockout OVCAR8 cells. (A)** Genome-wide Hi-C contact maps for two biological replicates of WT and ko2 cells. Blue arrows indicate some of representative multiple chromosomal rearrangements. **(B)** Spectral karyotype (SKY) analysis of OVCAR8 cells showing the complex aneuploid karyotype characteristic of this cell line. The representative SKY image was obtained from the NCBI SKY/M-FISH & CGH Database (PMID: 15934046). **(C)** Contact probability as a function of genomic distance (P(s)) for WT and ko2 Hi-C datasets at resolutions of 250, 100, 50, and 10 kb, demonstrating comparable distance-dependent contact decay. **(D)** Proportion of cis and trans chromatin interactions in WT and ko2 Hi-C libraries. **(E)** Fraction of long-range (>40 kb) cis interactions in two biological replicates of WT and ko2 cells. **(F)** Distribution of E1 compartment scores across the indicated genomic region (chr19:42,325,607-43,465,608), showing a significant increase in compartment strength in ko2 cells (Wilcoxon signed-rank test).

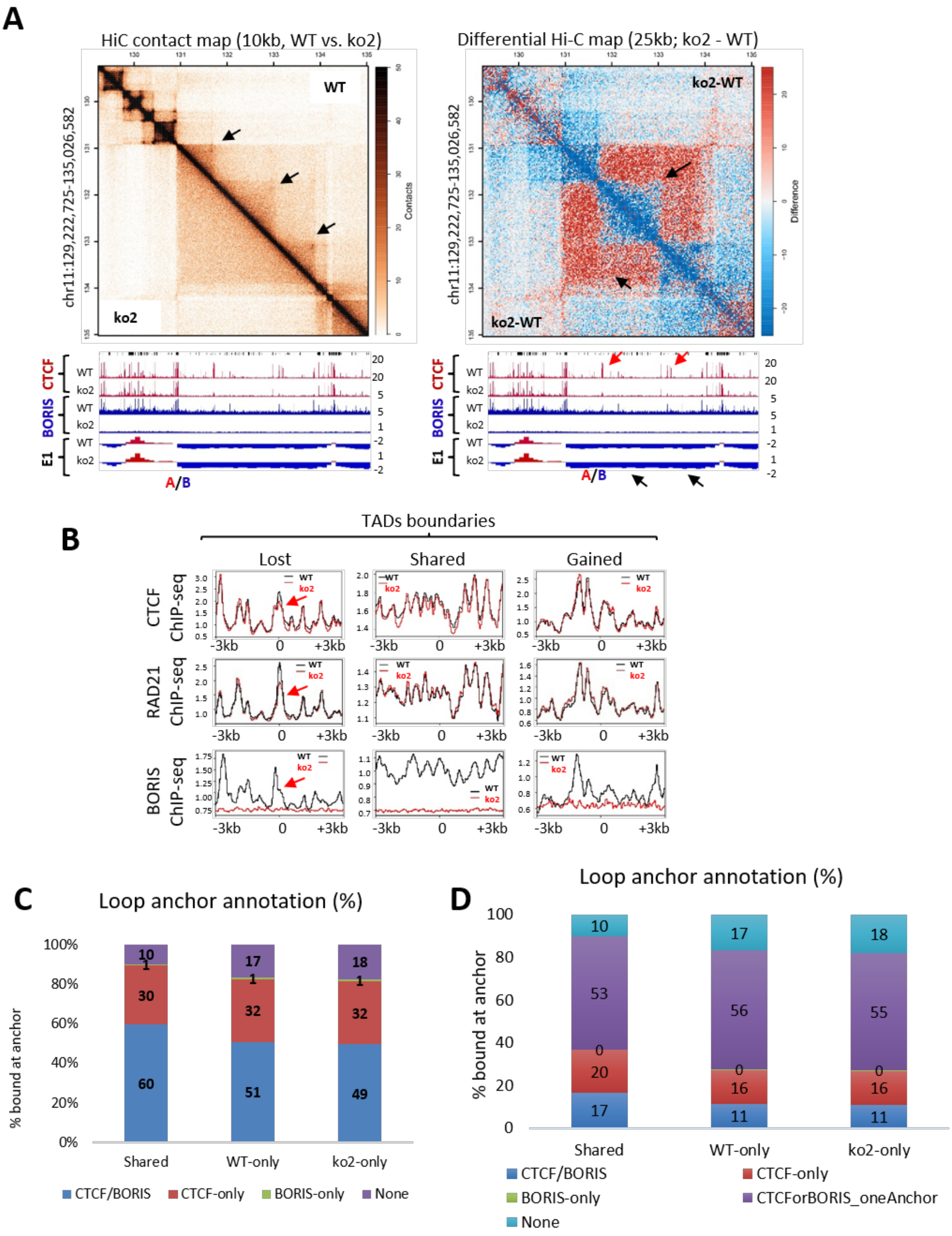
A
HiC contact map (10kb, WT vs. ko2)
Differential Hi-C map (25kb; ko2 - WT)
chr11:129,222,725-135,026,582
WT
ko2
ko2-WT
Contacts
Difference
CTCF
BORIS
E1
A/B
B
TADs boundaries
Lost
Shared
Gained
CTCF ChIP-seq
RAD21 ChIP-seq
BORIS ChIP-seq
-3kb
0
+3kb
C
Loop anchor annotation (%)
% bound at anchor
100%
80%
60%
40%
20%
0%
Shared
WT-only
ko2-only
CTCF/BORIS
CTCF-only
BORIS-only
None
D
Loop anchor annotation (%)
% bound at anchor
100
80
60
40
20
0
Shared
WT-only
ko2-only
CTCF/BORIS
CTCF-only
BORIS-only
CTCForBORIS_oneAnchor
None

**Figure S10. BORIS contributes to TAD boundary formation and chromatin loop organization in OVCAR8 cells. (A)** Representative Hi-C contact maps (left) and differential interaction map (right; ko2-WT) illustrating changes in chromatin interactions following BORIS knockout. Corresponding CTCF and BORIS ChIP-seq tracks and compartment score (E1) profiles are shown below. Arrows indicate representative changes in CTCF occupancy (red) and compartment scores (black) in ko2 cells compared to WT. **(B)** Average ChIP-seq signal profiles of CTCF, RAD21, and BORIS centered on TAD boundaries that are lost, shared, or gained following BORIS knockout (ko2) relative to wild-type (WT). Loss of CTCF and RAD21 occupancy at lost TAD boundaries (red arrows) coincides with loss of BORIS occupancy. Signals are displayed within ±3 kb of TAD boundaries. **(C-D)** Annotation of loop anchors shared between WT and ko2 or unique to either condition. Stacked bar plots show the proportion of loop anchors occupied by CTCF/BORIS, CTCF only, BORIS only, or neither factor at least one loop anchor **(C),** and the proportion of chromatin loops classified according to the occupancy of both loop anchors **(D).**
